# Cholangiocytic differentiation drives cell proliferation in hepatoblastoma through Wnt-dependent FGF19 signaling

**DOI:** 10.1101/2023.06.08.544267

**Authors:** Peng V. Wu, Matt Fish, Florette K. Hazard, Chunfang Zhu, Sujay Vennam, Hannah Walton, Maurizio Morri, Norma Neff, Robert B. West, Roel Nusse

## Abstract

Cancers evolve not only through the acquisition and clonal transmission of somatic mutations but also by non-genetic mechanisms that modify cell phenotype. Here, we describe how transcriptional heterogeneity arises within human hepatoblastoma, one of the cancers with the lowest mutational burden, characterized by activating mutations in the Wnt pathway. Histology-guided RNA sequencing and evaluation of spatial gene expression in primary hepatoblastomas identified foci of tumor cells within the highly proliferative embryonal histology that express the growth factor FGF19, colocalizing with markedly increased expression of Wnt target genes and cholangiocyte markers. In patient-derived tumoroids, FGF19 provided a required growth signal for FGF19-negative cells, and its expression depended on both Wnt/μ-catenin and the biliary transcription factor SOX4. Our results reveal that a biliary lineage program induces FGF19 as a paracrine signal for tumor growth, thereby modulating the transcriptional outcome of constitutive Wnt activation and tumor cell proliferation.

## INTRODUCTION

Cancers are classically thought to develop through clonal evolution, whereby sequential somatic mutations give rise to heterogeneous cell lineages subject to selective forces, leading to the emergence of subclones^1,2^. On the other hand, epigenetic mechanisms, which generate the vast heterogeneity of cells and tissues arising during normal embryogenesis, can significantly expand the diversity of cells carrying the same genetic mutations^3,4,5,6^. Such paths toward intra-tumoral heterogeneity may be particularly relevant for pediatric cancers with embryonic origins and relatively lower mutational burdens than adult cancers^7,8^. For instance, heterogeneity in neuroblastoma has been attributed to at least two differentiation states governed by distinct transcription factor networks^9,10^, and a transition from the nonadrenergic to the mesenchymal cell state leads to resistance to targeted immunotherapy^11^.

Hepatoblastoma is the most common pediatric liver malignancy, posited to originate from hepatoblasts, the embryonic progenitor of the two main cell types in the liver: hepatocytes and cholangiocytes. Several features render this tumor type an excellent model to study the relative contributions of genetic and non-genetic drivers of heterogeneity. Genomic studies have revealed a remarkably low tumor mutational burden, with up to 80-90% of hepatoblastomas containing mutations of the *CTNNB1* gene that abrogate degradation of the β-catenin protein, leading to constitutive activation of the Wnt pathway^12,13,14,15^. Despite the relative simplicity of their genomic alterations, hepatoblastomas nevertheless demonstrate significant phenotypic and histologic heterogeneity. Individual hepatoblastomas are often comprised of a mixture of epithelial histologies, known as fetal and embryonal, as well as areas of mesenchymal differentiation^16^. Tumors comprised purely of well-differentiated fetal histology exhibit low mitotic activity and are cured with surgical resection alone^17^, while those containing embryonal histology behave more aggressively. Thus, identifying biologic features that differentiate between the fetal and embryonal histologies has important treatment implications.

Previous studies using microarrays on bulk tumor tissue have ascribed differential Wnt-driven transcriptional programs to tumors comprised primarily of fetal histology as compared to those containing components of embryonal histology^18^, so called C1 vs. C2 tumors, respectively^19^. Further studies have proposed additional molecular classification schemes known as the C1/C2A/C2B^20^, MRS-1/MRS-2/MRS-3^21^, and the hepatocytic/liver progenitor/mesenchymal subgroups^22^, which stratify tumors into different prognostic categories. However, due to the rarity of this cancer and technical difficulties in propagating these cells *in vitro*, research into the functional relevance of specific genes and biomarkers has been hindered by the limited number of patient-derived cell lines^23^.

Although aberrant Wnt activation appears crucial to the development of hepatoblastoma, it remains unclear how heterogeneity in the expression of Wnt target genes arises and what accounts for the differences in proliferation between the fetal and embryonal histologies. Prior studies of hepatoblastoma cell lines have attributed roles for Wnt/β-catenin^19^, as well as the FGFs, FGF19 and FGF8, to cell proliferation^24,25^. Studies of normal hepatocytes suggest that Wnt and FGFs act at different steps in the cell cycle, as *in vitro* propagation of hepatocytes requires both exogenous Wnt agonists and growth factors acting through RTK/MAPK pathways^26,27^. In the normal liver, Wnt signals from endothelial cells in the central vein act on neighboring hepatocytes to promote mitosis^28^. Meanwhile, additional signals that drive hepatocyte cell cycle progression from G1 to S phase include the endocrine growth factor, FGF19^29^, normally expressed in the small intestine and gall bladder in response to bile acids^30^. In the embryonic liver, Wnt/β-catenin promotes hepatoblast maturation and survival^31^, while aberrant Wnt activation represses the hepatic fate and promotes biliary differentiation of hepatoblasts^32^. However, the implications of aberrant hepatobiliary differentiation for tumorigenesis in hepatoblastoma have not been reported.

Here, we use histology-guided RNA sequencing to elucidate the transcriptional heterogeneity within the fetal, embryonal, and mesenchymal components of hepatoblastoma. We uncover differentiation along the biliary and hepatic lineages as a key determinant of intra-tumoral heterogeneity and identify FGF19 as a context-specific target of Wnt that provides a requisite proliferative signal for surrounding tumor cells.

## RESULTS

### Histology-based RNA sequencing identifies increased expression of Wnt target genes, proliferative, and biliary markers in embryonal hepatoblastoma

To characterize the transcriptional profiles of distinct histologic components of hepatoblastoma, we used laser capture microdissection to isolate regions identified by a pathologist as embryonal, fetal, and mesenchymal histology, as well as adjacent normal liver, from a series of 17 patients with hepatoblastoma (Table S1). We performed RNA sequencing on 74 micro-dissected samples using a technique known as Smart-3SEQ^33^, which permits accurate quantification of transcript abundance from small, formalin-fixed, paraffin-embedded (FFPE) specimens in which RNA has degraded (Figure 1A, Table S2). In the initial quality control analysis, two samples were outliers, with low RNA yield and less than 100,000 uniquely mapped reads; these were excluded from the final analyses (Figure S1A). For the remaining 72 samples, the percentage of uniquely mapped reads was comparable to the range of 17-27% previously reported for Smart-3SEQ of FFPE specimens^33^ (Figure S1A).

**Figure 1.**
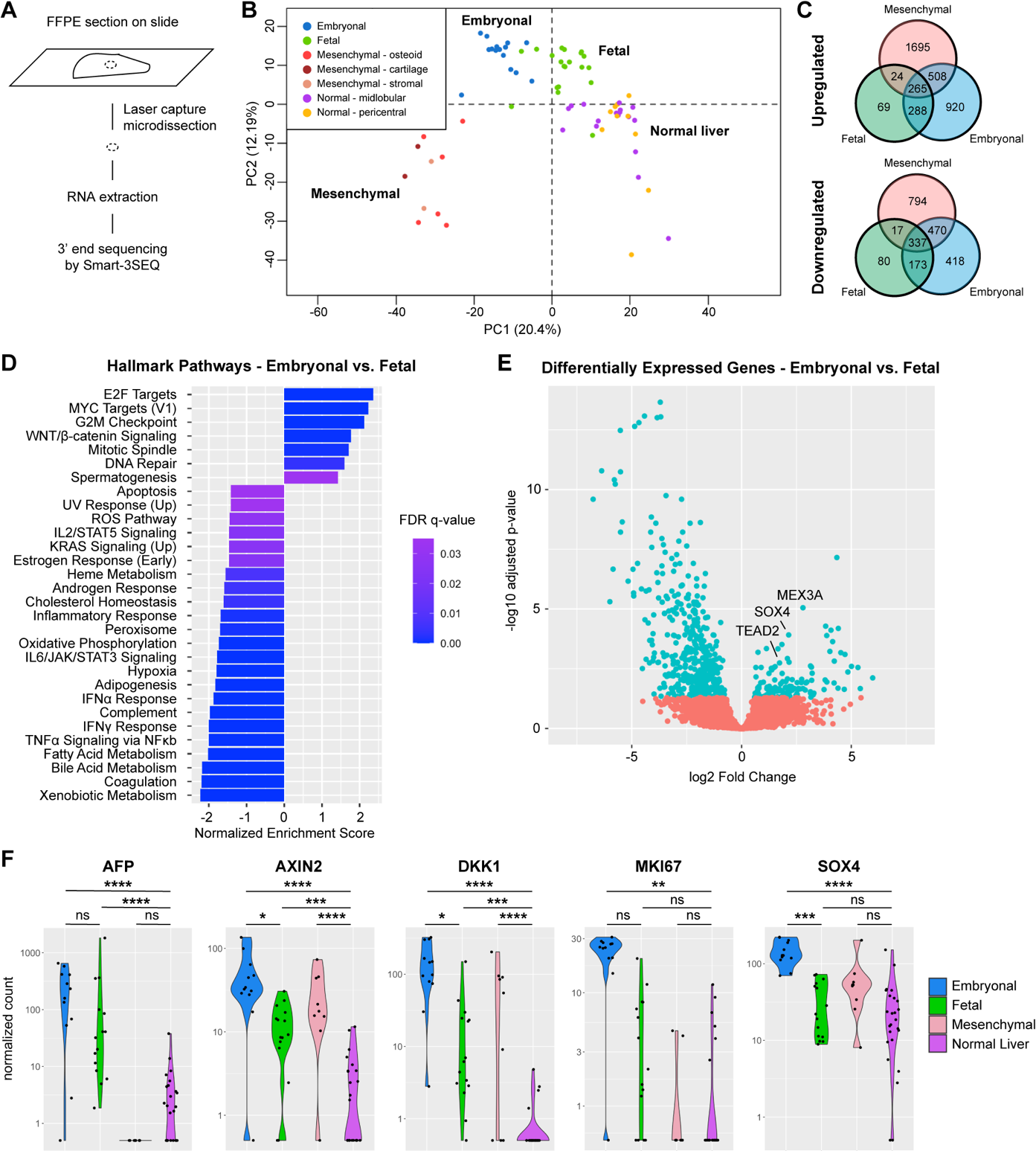
Histology-based RNA sequencing reveals increased expression of Wnt target genes, proliferation, and cholangiocyte markers in embryonal hepatoblastoma. A. Schematic of Smart-3SEQ. B. Principal component analysis plot of RNA sequencing from 72 samples with the histologies noted, using top 1500 expressed genes. C. Venn diagram of differentially expressed, upregulated and downregulated, genes (log2 fold change > 1, padj < 0.05) in embryonal, fetal, or mesenchymal hepatoblastoma compared to normal liver, determined by DESeq2. D. Top hallmark pathways differentially expressed between embryonal and fetal components of hepatoblastoma, using gene set enrichment analysis (GSEA) of lists ranked by p-value, obtained from DESeq2. E. Volcano plot of differentially expressed genes between embryonal and fetal components of hepatoblastoma with SOX4 and target genes labeled. F. Normalized counts of representative markers of Wnt pathway, proliferation, and cholangiocytic differentiation. * denotes padj <0.05, ** denotes padj <0.01, *** denotes padj <0.001, **** denotes padj <0.0001. ns = not significant. See also Figure S1.

Using unsupervised hierarchical clustering and principal component analysis to represent the transcriptional profiles of each sample in two dimensions, we found that the samples clustered based on their histologies rather than by patient or other patient-specific characteristics such as age, sex, or treatment history (Figure 1B, S1B). The transcriptional profiles of normal liver from either the pericentral or midlobular zones clustered together (Figure 1B), thus these samples were all included as normal liver in further analyses. Similarly, mesenchymal components dissected from areas that appeared to be osteoid, stromal, or cartilaginous clustered together (Figure 1B) and were collectively assigned as mesenchymal histology for additional analyses. Among the tumor components, the transcriptional profiles of specimens with fetal histology most resembled those of normal liver while those with mesenchymal histology diverged the most from normal liver (Figure 1B).

We next identified differentially expressed genes comparing the embryonal, fetal, and mesenchymal tumor components to normal liver, using DESeq2^34^ (Figure 1C, Table S3). Using a threshold of fold change > 2 and padj < 0.05 for statistical significance, we found that all the tumor components shared a common set of 265 upregulated and 337 downregulated genes (Figure 1C). The embryonal and mesenchymal components shared the most differentially expressed genes in common (508 upregulated, 470 downregulated), while the fetal and mesenchymal histologies shared the fewest differentially expressed genes (24 upregulated, 17 downregulated) (Figure 1C).

Gene set enrichment analysis (GSEA)^35,36^ revealed significant upregulation of Wnt/β-catenin signaling in all tumor components as compared to normal liver (Figure S1C). Both the fetal and embryonal components showed upregulation of E2F and MYC targets as well as G2M checkpoint components (Figure S1C). Unique to the embryonal component were DNA repair and mitotic spindle genes (Figure S1C). Meanwhile, the mesenchymal components showed upregulation of epithelial-mesenchymal transition and TGFβ signaling (Figure S1C). Comparing the embryonal to the fetal components of hepatoblastoma (Table S3), upregulated pathways included Wnt signaling, Myc targets, E2F targets, mitotic spindle, G2M checkpoint, and DNA repair genes (Figure 1D). The list of genes significantly upregulated in the embryonal compared to fetal components (Table S3) also included SOX4, a transcription factor important for biliary differentiation^37,38^, and two of its reported target genes, TEAD2^39^ and MEX3A^40^ (Figure 1E-F).

Focusing further on specific genes, we found that Wnt targets, such as AXIN2 and DKK1, were significantly upregulated in all components of the tumors, but particularly increased in the embryonal components (Figure 1F). Other known markers of hepatoblastoma, including AFP, DLK1, and GPC3, were upregulated in the epithelial (fetal and embryonal) but not mesenchymal components (Figure 1F, S1D). In addition to higher expression of Wnt target genes, embryonal components showed increased expression of markers of cell cycle progression, such as MKI67 and CCND2 (Figure 1F, S1D). Interestingly, as compared to normal liver, the embryonal components also showed higher expression of the biliary marker KRT19 and relatively lower expression of the hepatic marker HNF4A, while such differences were not observed in the fetal components (Figure S1D). Thus, the embryonal histology corresponded to higher expression of Wnt target genes, proliferation markers, and cholangiocytic markers, and we subsequently focused on understanding the relationship between these features.

### Focal FGF19 expression within highly proliferative regions of embryonal hepatoblastoma

Next, we examined the expression of candidate drivers of increased proliferation in embryonal hepatoblastoma. Interestingly, several FGFs, including FGF8, FGF9, and FGF19, were expressed at low levels in the embryonal components but not in the fetal components nor in normal liver (Figure 2A, S2A). In contrast, the growth factor IGF2, known to be upregulated in hepatoblastomas^41,42^, was similarly elevated in both the fetal and embryonal components (Figure S2A). Other growth factors expressed in normal liver, such as IGF1 and MET, the gene encoding for the hepatocyte growth factor, were not elevated in hepatoblastoma compared to normal liver, while EGF was not detected in the tumor regions or normal liver (Figure S2A).

**Figure 2.**
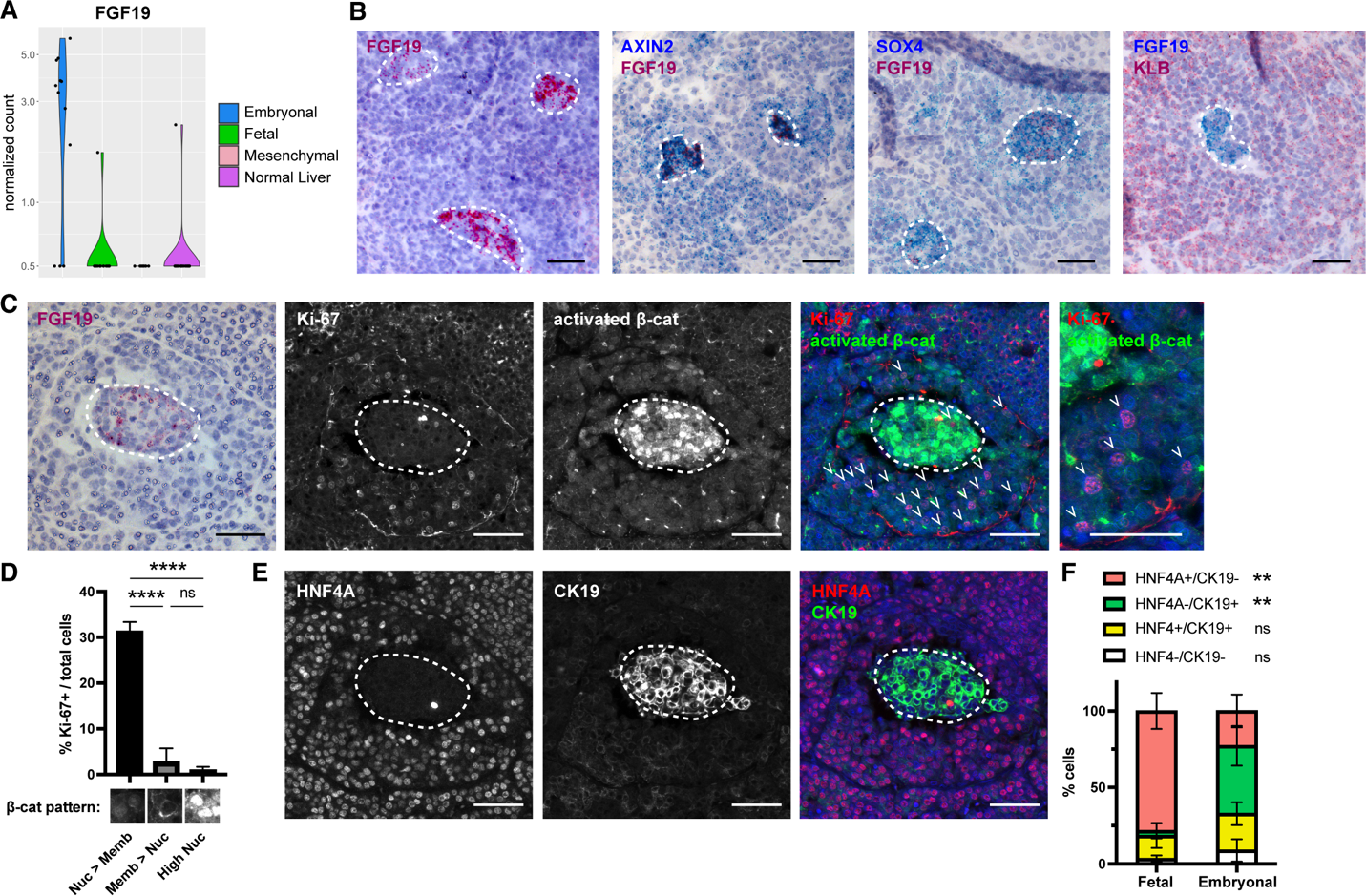
FGF19 is expressed focally within proliferative areas of embryonal hepatoblastoma. A. Normalized counts of FGF19 detected by Smart-3SEQ in the different histologic components of hepatoblastoma. B. RNA in situ hybridization in primary hepatoblastoma sections detecting the following genes - FGF19 (left); double in situ of AXIN2 (blue) and FGF19 (red) (panel 2 from left); double in situ of SOX4 (blue) and FGF19 (red) (panel 3 from left); double in situ of FGF19 (blue) and KLB (red) (panel 4 from left). White dashed lines outline cells expressing FGF19. Scale bar: 50 μm. C. Serial sections of primary hepatoblastoma showing RNA in situ hybridization of FGF19 (left), and co-immunofluorescence for Ki-67 (panel 2 from left) and activated unphosphorylated β-catenin (panel 3 from left), merged with DAPI (panel 4 from left), and at higher magnification (panel 5 from left). White dashed lines outline cells expressing FGF19. White arrowheads indicate Ki-67+ cells. Scale bar: 50 μm. D. Quantification of percentage of Ki-67+ cells with different β-catenin staining patterns as shown. (n = 3 primary hepatoblastoma specimens, >1000 cells scored). E. Co-immunofluorescence for HNF4A (left) and CK19 (middle), merged with DAPI (right) in serial sections of primary hepatoblastoma from C. White dashed line outlines cells expressing FGF19. Scale bar: 50 μm. F. Quantification of percentage of total cells with HNF4A^+^CK19^−^ (hepatocytic), HNF4A-CK19^+^ (cholangiocytic), HNF4A^+^CK19^+^ (hepatoblastic), and HNF4A^−^CK19^−^ (non-hepatobiliary) staining patterns in fetal and embryonal components of hepatoblastoma. (n = 4 primary hepatoblastoma specimens, >100 cells scored for each histology) ** denotes p<0.01 by student t-test, ns = not significant, comparing fetal vs. embryonal. See also Figure S2.

We were particularly interested in FGF19 as its secretion by a hepatoblastoma cell line has been previously reported to serve as an autocrine growth factor^24^. Furthermore, in mice, pericentral Wnt-responsive hepatocytes proliferate in response to FGF15, the mouse ortholog of FGF19, which is secreted by the small intestine^43^ during cycles of fasting and re-feeding and leads to cell proliferation in conjunction with Wnt signals^29^. We confirmed the presence of cells expressing FGF19 in primary hepatoblastoma specimens using RNA in situ hybridization (Figure 2B, summarized in Table S1). Strikingly, cells expressing FGF19 characteristically clustered within regions of embryonal histology with multiple FGF19-expressing foci appearing in the same tumor (Figure 2B). While the embryonal regions had higher AXIN2 and SOX4 expression compared to surrounding fetal regions, the FGF19-expressing foci showed particular upregulation of both AXIN2 and SOX4 (Figure 2B). To determine which cells might be responding to the FGF19 signal, we investigated the expression pattern of KLB, the co-receptor required for FGF19 signaling through FGFR4^44^. Unlike AXIN2 and SOX4, KLB did not co-localize with FGF19 but was highly expressed in cells surrounding the FGF19-expressing foci (Figure 2B), suggesting that FGF19 might serve as a paracrine signal.

We next interrogated the correlation of FGF19 expression in primary hepatoblastoma tissues with the proliferative index and pattern of β-catenin staining, where nuclear β-catenin indicates high levels of Wnt signaling, while normal hepatocytes exhibit primarily membrane-bound β-catenin. Immunofluorescence for activated (unphosphorylated) β-catenin in hepatoblastomas showed three patterns: cells with primarily membranous staining (Memb > Nuc in Figure 2D), those with primarily nuclear staining (Nuc > Memb in Figure 2D), and foci of cells with particularly high nuclear staining (High Nuc in Figure 2D) that co-localized with regions of FGF19 expression in serial sections (Figure 2C). Cells with primarily nuclear β-catenin staining (Nuc > Memb) surrounded the FGF19-expressing cells and were more likely to be positive for Ki-67 (marker of active cell cycling) and phospho-histone H3 (marker of mitosis) than either the FGF19-expressing cells with high nuclear β-catenin (High Nuc) or those with primarily membranous β-catenin staining (Memb > Nuc) (Figure 2C-D, S2B-C).

Given the correlation between expression of FGF19 and the biliary transcription factor SOX4, we investigated the differentiation state of these cells by co-immunofluorescence for the hepatic marker HNF4A and the biliary marker CK19 (Figure 2E). We found that foci of FGF19 expression co-localized with cells that were HNF4A^−^ CK19^+^, which we denote as cholangiocytic. Upon quantification, such cholangiocytic cells were more frequently found in embryonal as compared to fetal regions of hepatoblastoma (Figure 2F). Meanwhile, in the embryonal regions, there were significantly less HNF4A^+^CK19^−^ (hepatocytic) cells, while both fetal and embryonal components contained similar numbers of HNF4A^+^CK19^+^ (hepatoblastic) cells and low numbers of HNF4A^−^CK19^−^ (non-hepatobiliary) cells (Figure 2E-F). Co-localization of CK19 and Ki-67 showed a high proliferative index among hepatoblastic cells compared to low (but not absent) proliferation of the cholangiocytic and hepatocytic cells (Figure S2D). Thus, FGF19 expression correlates with a cholangiocytic differentiation state.

### Patient-derived hepatoblastoma tumoroids recapitulate embryonal and fetal gene signatures

To develop an experimental system to test the functional requirements of the different signaling pathways in hepatoblastoma, we obtained fresh tumor specimens from patients and generated primary cultures using our recently published protocol to propagate hepatoblastoma cells as 3D tumoroids^45^. Cell clusters were embedded in Matrigel and propagated in media containing the growth factors, EGF, FGF10, and HGF. Of note, the media did not include exogenous Wnt agonists, forskolin, or TNF-α, which have previously been used in the culture of normal hepatocytes^26,27^. In fact, the addition of the GSK3β inhibitor, CHIR99021, a small molecule Wnt pathway agonist, did not enhance the establishment of the tumoroid cultures (data not shown).

From a series of 16 patients, we successfully obtained 10 independent cultures that could be serially passaged as tumoroids for >100 days (Table S4). Morphologically, the tumoroids formed spherical cell clusters or occasionally multilobulated structures (Figure 3A). The patients from whom these tumoroids derived were representative of the general population of patients with hepatoblastoma, with a median age of 19 months (range: 4-129 months) and a slight male predominance (62%). Four of the tumoroids were obtained prior to any chemotherapy, while the remainder were isolated at the time of tumor resection following neoadjuvant chemotherapy (Table S4).

**Figure 3.**
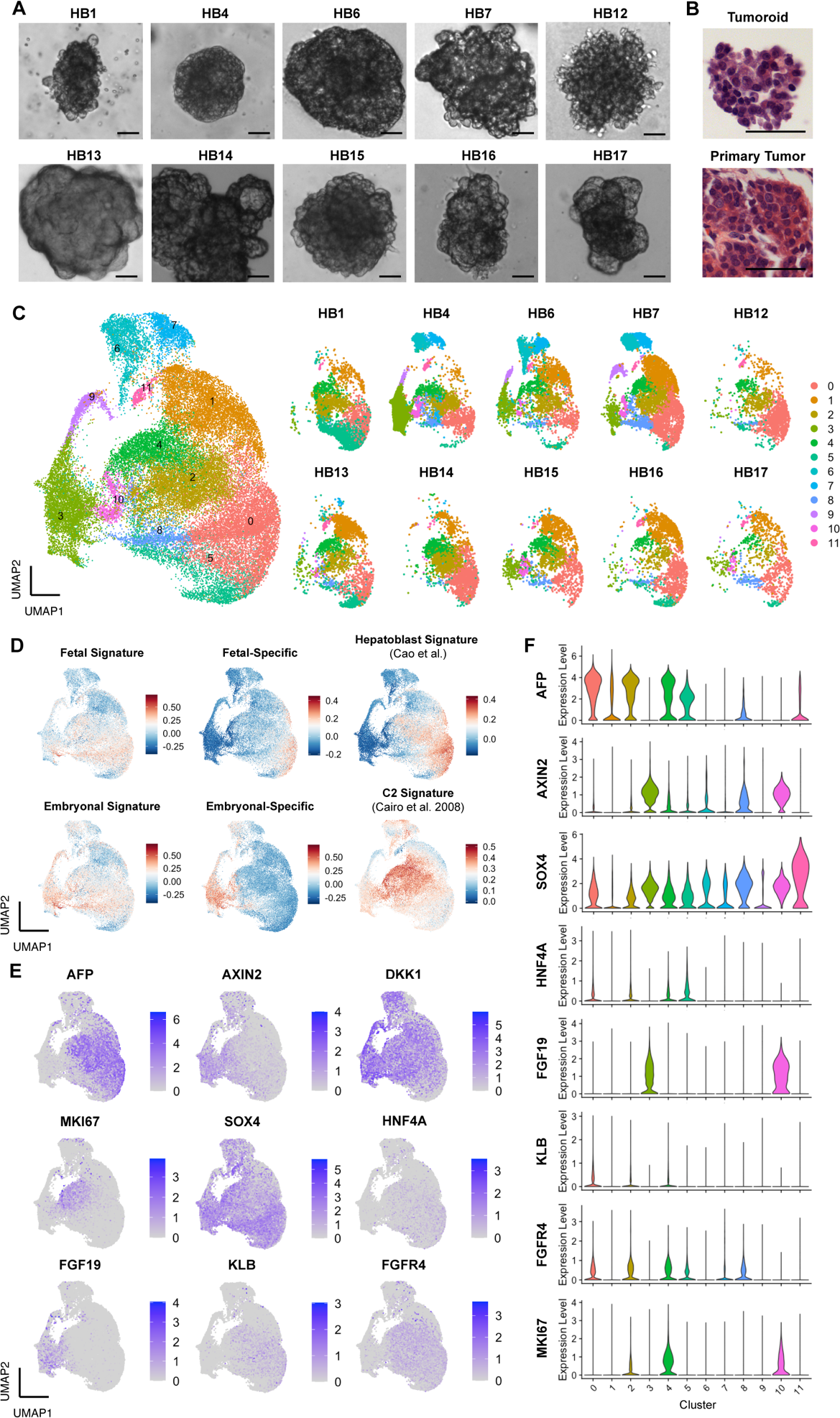
Hepatoblastoma tumoroids recapitulate expression of embryonal and fetal gene signatures. A. Representative brightfield images of primary hepatoblastoma tumoroids at early passage (P1-2) from patients indicated. Scale bar: 50 μm. B. Representative H&E image of early passage tumoroid compared to the primary tumor from the same patient. Scale bar: 50 μm. C. UMAP representation of scRNA sequencing results of 10 patients, colored by cluster, along with separate UMAP plots for each patient. F. UMAP plots showing relative expression of different gene signatures defined as follows and listed in Table S6: fetal = top 50 differentially upregulated genes between the fetal histology vs. normal liver; embryonal = top 50 differentially upregulated genes between the embryonal histology vs. normal liver; fetal-specific = top 50 differentially downregulated genes between the embryonal vs. fetal histologies; embryonal-specific = top 50 differentially upregulated and downregulated genes between the embryonal vs. fetal histologies; normal hepatoblast signature^47^; and C2 hepatoblastoma signature^19^. E. UMAP plots showing relative gene expression for hepatoblastoma markers, Wnt target genes, proliferation and differentiation markers, and FGF19 and its receptor/co-receptor, FGFR4/KLB. F. Violin plots showing expression levels of marker genes by cluster. See also Figure S3.

We first investigated the status of the *CTNNB1* gene in the tumoroids, since activating Wnt pathway mutations have been described in the majority of hepatoblastomas^12,13,14,15^ and all the tumoroids grew in media without an exogenous Wnt agonist. We found evidence of mutations involving exon 3 of *CTNNB1* in all of the tumoroids, ranging from deletion of the entire exon to point mutations in residues important for GSK3β phosphorylation and subsequent targeting of β-catenin for ubiquitin-mediated degradation (Table S4). Targeted cancer gene panel sequencing of the primary tumor specimens identified few additional mutations, including one patient each with pathogenic germline mutations in *APC* and *ARID1A*, while the remaining mutations identified were variants of unclear significance (Table S4).

To determine whether the tumoroids captured the transcriptional heterogeneity of primary hepatoblastomas, we queried gene expression at early passage using single cell RNA sequencing. We analyzed gene expression data for a total of 40,666 cells from tumoroids corresponding to the 10 different patients and clustered the cells using Seurat and UMAP visualization^46^ (Figure 3C, Table S5). Clusters 0, 2, and 4, characterized by high expression of known hepatoblastoma markers, including AFP, DLK1, and GPC3, were shared among all the tumoroids, while the remaining clusters of cells were more heterogeneously represented across the different tumoroids (Figure 3C, Figure 3E-F, Figure S3B, Table S5).

We next determined whether the transcriptional profiles of the hepatoblastoma cells *in vitro* recapitulated the histology-based transcriptional signatures identified in the primary tumors. We used the top 50 genes (ranked by p-value) that were differentially upregulated in the fetal or embryonal components of hepatoblastoma compared to normal liver, as determined by Smart-3SEQ, to calculate the fetal or embryonal signature, respectively, for each cell (Figure 3D, Table S6). As there was significant overlap in cells expressing both fetal and embryonal signatures (in clusters 0, 2, 4, and 8), we next used the top 50 genes differentially upregulated or downregulated in the fetal compared to embryonal components to identify clusters expressing a fetal-specific or embryonal-specific gene signature (Figure 3D, Table S6). While the fetal-specific signature was highest in a subset of cell clusters 0 and 1, the embryonal-specific signature was highest in clusters 3, 6, 8, 9, and 10 (Figure 3D).

Finally, we calculated a mesenchymal signature for each cell based on the top 50 genes differentially upregulated in the mesenchymal components compared to normal liver (Figure S3A, Table S6). Although there was significant overlap among cells expressing the mesenchymal signature with those expressing the embryonal-specific signature (Figure S3A, Figure 3D), almost no cells showed high expression of a mesenchymal-specific signature, derived from genes differentially upregulated in the mesenchymal compared to the embryonal components of the hepatoblastomas (Figure S3A, Table S6). Overall, the single cell expression data as well as the histologic appearance of the tumoroids (Figure 3B) suggested that the cultured cells represented the epithelial components of primary hepatoblastomas, encompassing cells with both fetal and embryonal patterns of gene expression.

Comparing to published studies, we found that cells expressing either the fetal or embryonal signatures overlapped with cells highly expressing upregulated genes previously reported in hepatoblastoma^19^ (Figure S3A). Meanwhile, cells with a higher fetal-specific signature correlated with higher expression of genes previously reported to be upregulated in normal fetal hepatoblasts^47^ (Figure 3D), while the cells with the embryonal-specific signature comprised a subset of cells with high expression of the C2 gene signature, previously attributed by microarray studies to embryonal hepatoblastoma and correlating with a worse outcome^19^ (Figure 3D).

We next analyzed each cluster individually, focusing on differences between cells expressing the embryonal-specific and those expressing the fetal-specific signature. In particular, the embryonal-specific clusters 3 and 10 were marked by upregulation of FGF19, as well as AXIN2 and SOX4 (Figure 3E-F, Figure S3C, Table S5), analogous to the FGF19-expressing cells that we identified in the primary hepatoblastomas (Figure 2B). These cells also showed high expression of FGF3, FGF8, and FGF9 (Figure S3B-C). Cycling tumor cells, marked by MKI67, primarily belonged to clusters 2, 4, and 10 (Figure 3E-F, Figure S3C), which also correlated with high expression of the C2 signature (Figure 3D). A subset of the MKI67-expressing cells in clusters 2 and 4 overlapped with the expression of the receptor/co-receptor FGFR4/KLB (Figure 3E-F, Figure S3C). This pattern of high FGF19 expression in a population of cells distinct from those expressing the cognate receptors resembled the expression pattern in primary tumors (Figure 2B), again suggesting that FGF19 may promote proliferation in a paracrine fashion.

### Patient-derived hepatoblastoma cells require growth factors for proliferation

Since the hepatoblastoma tumoroids grew in serum-free media containing well-defined chemical components and recombinant growth factors, we could systematically test the requirements for exogenous growth factors. For the majority (67%) of tumoroids tested, colonies could not form from single cells when grown in media lacking the standard growth factors, EGF, FGF10, and HGF (Figure 4A-B). Each of the growth factors alone stimulated colony formation to varying degrees but were redundant to the other two factors (Figure S4A-B). In contrast, this dependence on exogenous growth factors did not apply to the HepG2 cell line, which has an activating NRAS mutation^48^ and formed colonies even in the absence of EGF, FGF10, or HGF (data not shown).

**Figure 4.**
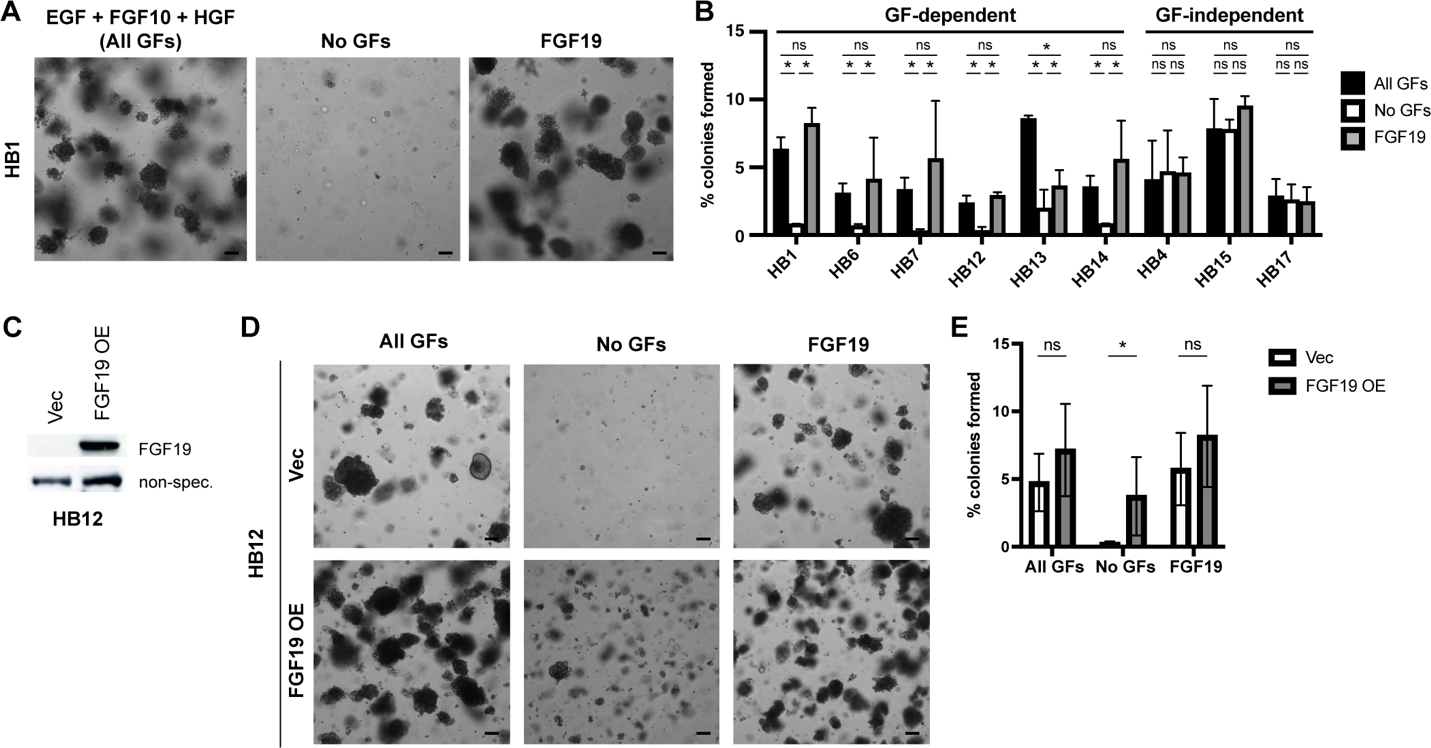
FGF19 is sufficient for hepatoblastoma colony formation *in vitro*. A. Brightfield images of hepatoblastoma colonies from HB1 after seeding as single cells and growing for 2 weeks in media containing EGF 50 ng/ml + FGF10 100 ng/ml + HGF 25ng/ml (All GFs), no GFs, or in media with FGF19 100 ng/ml. Scale bar: 100 μm. B. Quantification of colony assay in media with all GFs, no GFs, or FGF19 for each of 9 tumoroids. C. Immunoblot detection of FGF19 and a control non-specific band in HB12 cells transduced with lentiviral vector control or FGF19. D. Brightfield images of HB12 colonies transduced with lentiviral vector control or FGF19, seeded after 3 days of antibiotic selection as single cells, and grown for 2 weeks in media containing EGF + FGF10 + HGF (All GFs), no GFs, or in media with FGF19. Scale bar: 100 μm. E. Quantification of colony assay for HB12 as in D (n = 3 independent experiments, error bars indicate standard deviation. * denotes p<0.05 by paired student t-test). See also Figure S4.

We next tested whether FGF19, one of the FGF family members expressed in the primary hepatoblastomas (Figure 2A-B), could substitute by itself for the traditional growth factors used in the media. Indeed, for each of the tumoroids that required exogenous growth factors for colony formation, FGF19, when added to the basal media, was sufficient to promote colony formation (Figure 4A-B). In addition, overexpression of FGF19 in HB12, one of the tumoroids with low FGF expression at baseline (Figure S3D-F), permitted colony formation in the absence of growth factors (Figure 4C-E).

We further investigated the downstream mechanism by which these exogenous growth factors exert their effects on cell proliferation. Inhibition of the MAP kinase pathway using the MEK inhibitor U0126 reduced colony formation in all the tumoroids except HB13, which was nevertheless sensitive at higher concentrations (Figure S4C-D). A second MEK inhibitor, trametinib, also abrogated colony formation, with higher potency than U0126 (Figure S4E). Finally, GF-dependent tumoroids treated with MEK inhibition and withdrawal of exogenous growth factors arrested predominantly in G0/G1 phase of the cell cycle (Figure S4F-G). When the MEK inhibitor was washed out and media containing growth factors was added to the tumoroids, the cells progressed into S phase with 37 ± 8% EdU incorporation, as compared to 18 ± 2% when grown in media without growth factors (Figure S4F-H). These results suggested that despite constitutive Wnt pathway activation, the tumor cells require additional growth factors acting through the MAPK pathway to progress from G1 into S phase.

### FGF19 expression bypasses the requirement for exogenous growth factors

We next sought to understand how the growth factor-independent tumoroids (HB4, HB15, and HB17) could form colonies even in the absence of exogenous growth factors. These tumoroids showed a high percentage of cells expressing FGFs, particularly FGF19, by single cell RNA sequencing (Figure S3D-E). To test whether these tumor-intrinsic growth factors could provide a paracrine effect, we investigated whether conditioned media from HB4 and HB15, growing in the absence of exogenous growth factors, could support the colony formation of GF-dependent tumoroid cells. Indeed, HB1 cells, which require exogenous growth factors, formed colonies when grown in conditioned media from either HB4 or HB15 (Figure 5A-B). Colony formation in these conditions was inhibited by the MEK inhibitor, U0126 (Figure 5A-B), consistent with the hypothesis that the growth factor(s) secreted by HB4 and HB15 activate MAPK signaling in HB1 cells.

**Figure 5.**
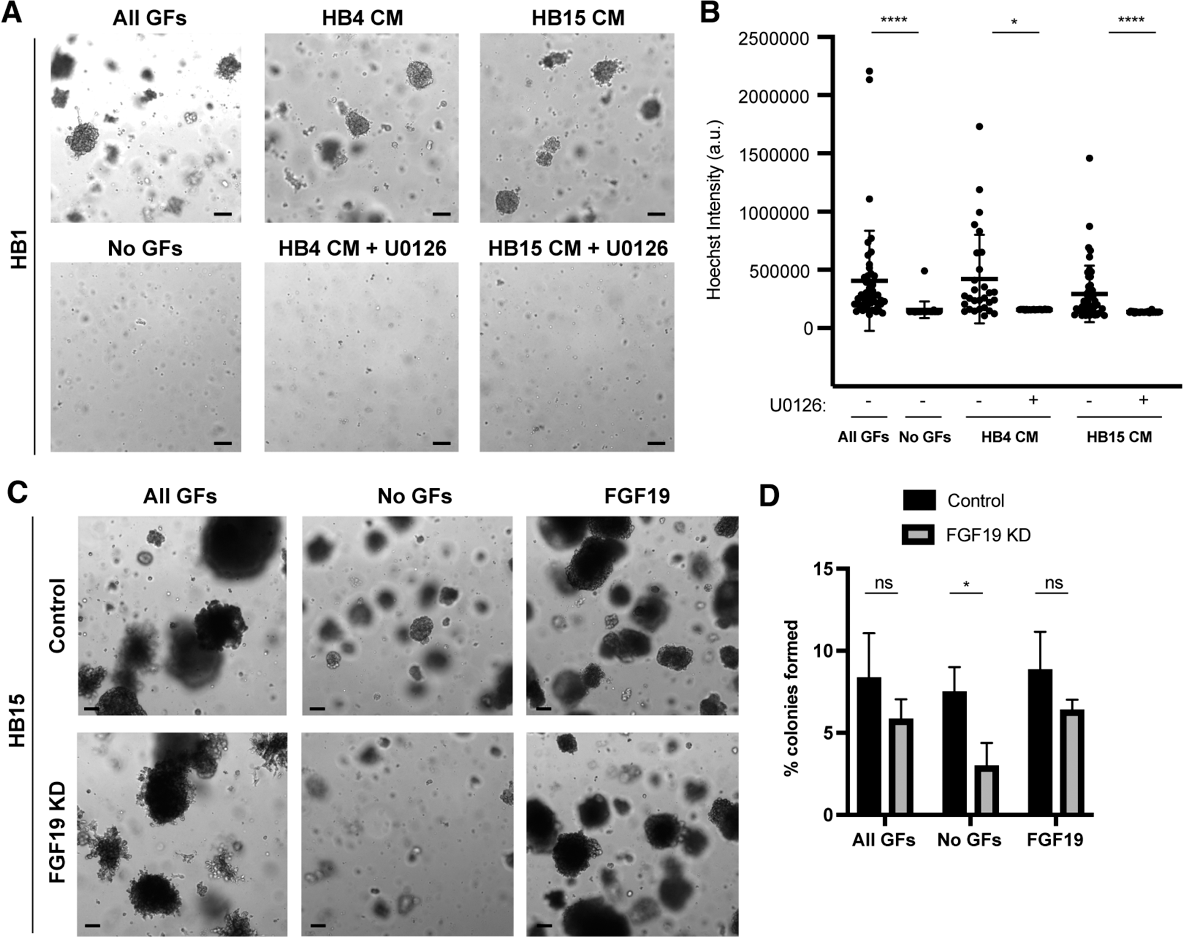
FGF expression bypasses the requirement for exogenous growth factors. A. Brightfield images of HB1 seeded as single cells and grown for 2 weeks in media containing EGF + FGF10 + HGF (All GFs), no GFs, HB4 conditioned media (CM) without or with U0126 (5 μM), and HB15 conditioned media (CM) without or with U0126 (5 μM). Scale bar: 100 μm. B. Quantification of Hoechst intensities of tumoroid colonies from A (**** denotes p<0.0001, * denotes p<0.05 by Kruskal-Wallis with Dunn’s multiple comparisons test). C. Brightfield images of HB15 colonies transduced with lentiviral shRNA to luciferase (control) or FGF19 seeded as single cells after 3 days of antibiotic selection and grown for 2 weeks in media containing EGF + FGF10 + HGF (All GFs), no GFs, or in media with FGF19. Scale bar: 100 μm. D. Quantification of colony assay as in B (n = 3 independent experiments, error bars indicate standard deviation. * denotes p<0.05 by paired student t-test). See also Figure S5.

To determine whether expression of FGF19 by a subpopulation of tumoroid cells was required for independence from exogenous growth factors, we evaluated colony formation upon inhibition of FGF19 signaling. Indeed, depleting FGF19 in HB15 using a lentiviral shRNA (Figure S5A) reduced the ability to form colonies in the absence of exogenous growth factors (Figure 5C-D). This was rescued by the presence of either exogenous FGF19 or the combination of EGF, FGF10, and HGF (Figure 5C-D). FGF19 knockdown in both HB4 and HB17 similarly reduced colony formation in the absence exogenous growth factors (Figure S5B). Consistent with the requirement for FGF19 signaling, the addition of BLU9931, a selective FGFR4 inhibitor^49^, to the GF-independent tumoroid cells also reduced colony formation in the absence of exogenous growth factors, with an average IC50 of 1.24 ± 0.31 μM across the three different tumoroids (Figure S5C). Thus, while these hepatoblastoma tumoroids did not require exogenous growth factors, they depended on their own expression of FGF19 and signaling through FGFR4.

### FGF19 expression depends on Wnt/β-catenin and biliary transcription factor SOX4

Next, we investigated the mechanism by which FGF19 is induced in hepatoblastoma cells. In normal tissues that express FGF19, transcriptional regulation occurs through the farnesoid X receptor (FXR) and FXR response elements, which are activated by bile acids^50,43,51^. In HB15 cells, FGF19 expression was not increased by the addition of bile acids (cholic acid and chenodeoxycholic acid) or the FXR agonist cilofexor, nor reduced by an FXR antagonist, guggulsterone^52^, suggesting that the regulation of FGF19 in these cells is uncoupled from FXR (Figure S6A-D).

Since FGF19 expression colocalized with higher Wnt target gene expression in the embryonal components of hepatoblastoma (Figure 2B), we asked whether the upregulation of FGFs in these tumoroids depended on β-catenin. In HB15, both lentiviral shRNA knockdown of CTNNB1 and overexpression of a dominant negative mutant of the transcription factor TCF4 (DN TCF4), which blocks β-catenin-mediated transcription^53^, resulted in >2-fold reduction in the expression of FGF19, FGF8, and FGF9 (Figure 6A). Consistent with the requirement of FGFs for proliferation of these cells, inhibition of β-catenin mediated transcription also reduced colony formation in the absence of exogenous growth factors (Figure 6B-C, Figure S6E-F). Colony formation was only partially rescued by addition of recombinant FGF19 (Figure 6B-C, S6E-F), suggesting that in the absence of active Wnt signaling, FGF19 alone is insufficient to drive cell proliferation. Instead, the combination of constitutive Wnt pathway activation and Wnt/β-catenin-dependent FGF19 together allow these tumor cells to progress through the cell cycle and form colonies in the absence of exogenous growth factors.

**Figure 6.**
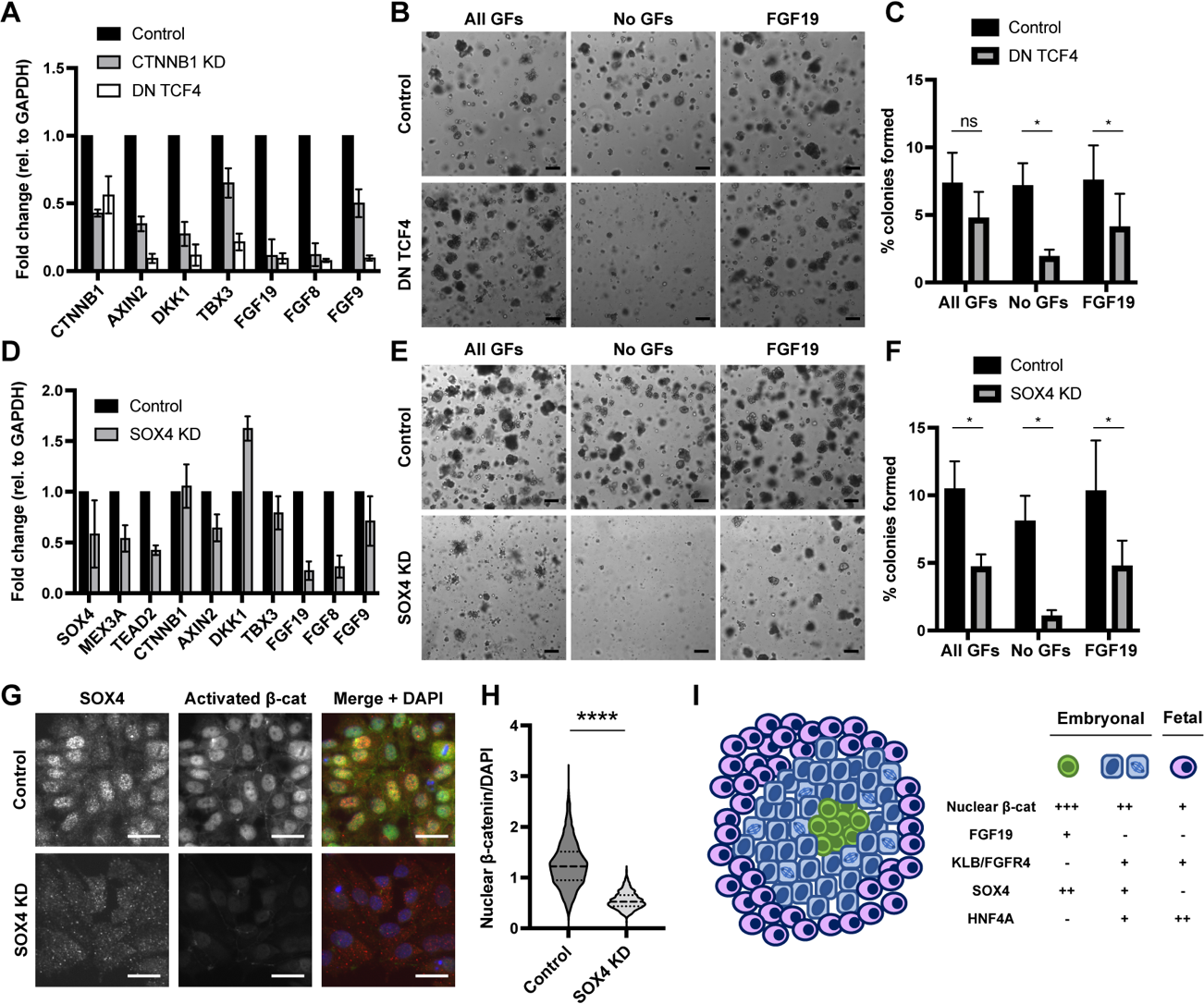
FGF19 expression depends on β-catenin and SOX4. A. qRT-PCR for Wnt target genes, FGF19, FGF8, and FGF9 in HB15 cells transduced with lentiviral shRNA to luciferase (control) or CTNNB1, or overexpressing dominant negative TCF4, assayed after 3 days of antibiotic selection. B. Brightfield images of HB15 colonies transduced with lentiviral GFP or DN TCF4, seeded after 3 days of antibiotic selection as single cells, and grown for 2 weeks in media containing EGF + FGF10 + HGF (All GFs), no GFs, or in media with FGF19. Scale bar: 200 μm. C. Quantification of colony assay as in B (n = 3 independent experiments, error bars indicate standard deviation. * denotes p<0.05 by paired student t-test. ns = not significant). D. qRT-PCR for SOX4 targets, Wnt target genes, FGF19, FGF8, and FGF9 in HB15 cells transduced with a control or SOX4 lentiviral shRNA, assayed after 3 days of antibiotic selection. E. Brightfield images of HB15 colonies transduced with lentiviral shRNA to luciferase (control) or SOX4, seeded after 3 days of antibiotic selection as single cells, and grown for 2 weeks in media containing EGF + FGF10 + HGF (All GFs), no GFs, or in media with FGF19. Scale bar: 200 μm. F. Quantification of colony assay as in E (n = 3 independent experiments, error bars indicate standard deviation. * denotes p<0.05 by paired student t-test). G. Co-immunofluorescence of SOX4 and activated β-catenin in HB15 cells transduced with lentiviral shRNA to luciferase (control) or SOX4. Scale bar: 25 μm H. Quantification of nuclear β-catenin intensity normalized to DAPI in HB15 cells transduced with lentiviral shRNA to luciferase (control) or SOX4, as in G (**** denotes p<0.0001 by Mann-Whitney test). I. Model depicting spatial organization of different tumor cell types in hepatoblastoma with table showing relative levels of nuclear β-catenin, FGF19, KLB/FGFR4, SOX4, and HNF4A. See also Figure S6.

Finally, we determined whether FGF19 expression in HB15 cells depends on the biliary transcriptional program, as FGF19 expression had colocalized with high SOX4 and CK19 in primary hepatoblastomas (Figure 2B). Depletion of SOX4 by lentiviral shRNA in HB15 reduced the expression of known SOX4 targets, such as MEX3A and TEAD2, while canonical Wnt targets, such as AXIN2 and DKK1, were only modestly affected (Figure 6D). Meanwhile, SOX4 knockdown significantly reduced the expression of FGF19, FGF8, and to a lesser extent FGF9 (Figure 6D). SOX4 depletion also reduced colony formation both in the absence and presence of growth factors, suggesting that SOX4 promotes colony formation through induction of FGF expression as well as an FGF-independent mechanism (Figure 6E-F). Although SOX4 knockdown did not affect expression of CTNNB1, nuclear β-catenin was substantially reduced (Figure 6G-H). Consistent with SOX4 acting upstream of Wnt/β-catenin, depletion of CTNNB1 did not significant affect the expression of SOX4 or its targets (Figure S6G). These results suggest that SOX4, in addition to driving the biliary transcriptional program, modulates the transcriptional output of a subset of β-catenin targets, including FGF19.

## DISCUSSION

While nearly all hepatoblastomas contain somatic activating mutations in *CTNNB1*, here we find evidence that the transcriptional output of Wnt pathway activation is modulated within individual tumors and exists along a gradient across the different histologies, with implications for their biologic behavior. We confirm lower expression of Wnt target genes and lower levels of nuclear β-catenin within fetal regions of hepatoblastoma compared to the embryonal regions, as has been previously reported^19,54^. Within the embryonal histology, we find regions of particularly high Wnt target gene expression and high nuclear β-catenin levels corresponding to high expression of cholangiocyte markers and a low proliferative index, surrounded by a zone of intermediate Wnt target gene expression and nuclear β-catenin, with decreased cholangiocytic markers and increased proliferation. In SOX4-expressing cholangiocytic tumor cells, higher nuclear β-catenin levels result in expression of FGF19, which promotes proliferation of surrounding HNF4A-expressing hepatoblastic/hepatocytic cells (Figure 6I).

We have thus uncovered a tumor-intrinsic source of growth factors that leads to increased proliferation within the embryonal components of hepatoblastoma. Although the expression of FGF19 and FGF8 have previously been reported in hepatoblastoma cell lines and primary tumors^24,25^, our study now ascribes their expression to a subset of cells with embryonal histology that show cholangiocytic differentiation. Importantly, we have localized FGF-expressing cells spatially within primary hepatoblastoma tissues, where they form distinctive foci in regions of high proliferative activity, resembling morphogen-secreting signaling centers in embryonic development. Our results indicate that activated Wnt/β-catenin is insufficient for tumor cell cycle progression and that FGFs or other growth factors are required to promote the transition from G1 to S phase. Of note, genomic amplification of *FGF19* has previously been identified in hepatocellular carcinoma^55,56^ and more rarely in hepatoblastoma^22^, supporting the notion that FGF19 signaling can provide a growth advantage in liver cancers.

Our finding that hepatoblastoma cells require exogenous growth factors despite constitutive activation of the Wnt pathway indicates that these cells are not intrinsincally self-sufficient. Although embryonal hepatoblastoma cells can bypass the requirement for exogenous growth factors through the expression of FGFs, including FGF19, our studies suggest that a significant portion of hepatoblastoma cells do not proliferate in the absence of growth factors from cell-extrinsic sources. These findings have significant implications for tumorigenesis in infancy and childhood, as hormones and other signals that promote organismal growth may also influence the proliferation of incipient cancer cells. Given that FGF19 is secreted by the intestine in response to feeding and bile acid secretion^30^, our studies warrant further investigation as to how endocrine signals and environmental modifications such as diet may promote the growth of hepatoblastoma. Our newly derived compendium of primary hepatoblastoma tumoroids provides a powerful platform to further elucidate the effects of different growth factors and nutrient-signaling pathways.

Overall, our findings reveal a non-genetic mechanism by which tumor cell proliferation is modulated by the developmental lineage programs of the hepatobiliary progenitors from which hepatoblastoma derives. Rather than simply a bystander effect, lineage-driven heterogeneity in the transcriptional outcomes of activated Wnt signaling results in a functional growth advantage for the bulk tumor. Our results support a cooperative model of tumor development whereby cholangiocytic tumor cells promote the proliferation of surrounding tumor cells expressing hepatic markers, linking cell state and function within the tumors. The relative tumor initiating capacity of these different cancer cell types individually or in combination remains to be tested and likely depends not only on their ability to respond to proliferative signals but also on the level of plasticity between the cell states.

## Supporting information

Supplemental Information including Tables and Figures

## ACKNOWLEDGEMENTS

We are grateful to the patients and families who consented to banking tissue for research. We thank the Bass Center Tissue Bank and members of the Pediatric Hematology/Oncology, Liver Transplant Surgery, Pediatric Interventional Radiology, and Pathology teams at Lucile Packard Children’s Hospital/Stanford Health Care for assistance with tissue procurement. We acknowledge the Stanford Functional Genomics Facility and the Chan Zuckerberg Biohub for sequencing support. We further thank the following individuals: Shirley Kwok and Pauline Chu from the Stanford Department of Pathology for tissue sectioning and slide preparation; Joanna Przybyl for instruction on analysis of RNA sequencing data; D. Berfin Azizoglu, Catriona Logan, and Xin Wang for valuable comments on the manuscript; and members of the Nusse lab for scientific and technical advice. This work was funded by the Howard Hughes Medical Institute (R.N.), the Stanford Department of Pathology (R.B.W.), the Stanford Maternal and Child Health Institute (Ernest and Amelia Gallo Clinical Fellowship to P.V.W.), and the Damon Runyon Cancer Research Foundation (Damon Runyon-Sohn Pediatric Cancer Fellowship Award, DRSG-28P-19, to P.V.W.).

## AUTHOR CONTRIBUTIONS

P.V.W. and R.N. conceived the project, designed experiments, supervised work, and wrote/edited the original manuscript. P.V.W. performed all experiments except the following: M.F. performed RNAscope in situ hybridization, F.K.H. assigned pathology to hepatoblastoma specimens, C.Z. prepared Smart-3SEQ libraries from micro-dissected samples, S.V. performed processing and alignment of Smart-3SEQ data, and H.W. performed and quantitated co-immunofluorescence of phospho-histone H3 and activated β-catenin in hepatoblastoma sections. P.V.W. performed all other data curation, formal analyses, programming, and code implementation. M.M. and N.N. provided resources for scRNA sequencing. R.B.W. provided resources for and supervised the Smart-3SEQ experiments.

## RESOURCE AVAILABILITY

Single cell RNA sequencing data have been uploaded to the Gene Expression Omnibus (GEO) and will be available upon publication. Hepatoblastoma tumoroids will be available upon request after publication. Further information and requests for resources and reagents should be directed to and will be fulfilled by the lead contact, Peng V. Wu.

## METHODS

### Human tissues

All human subjects research at Stanford University was approved by the Institutional Review Board. Archival hepatoblastoma specimens from 2003-2017 were identified through the Stanford Department of Pathology. Human hepatoblastoma specimens from patients diagnosed at Lucile Packard Children’s Hospital between 2018-2023 were collected at the time of biopsy or resection and processed as described below. Informed consent was obtained from the parents of all patients from whom fresh specimens were collected. All patient-derived tissues and cells described in this study were obtained at Stanford/Lucile Packard Children’s Hospital except for HB7, which was a gift from Bruce Wang (UCSF)^57^.

### Laser capture microdissection & Smart-3SEQ

Hematoxylin and eosin-stained sections of archival hepatoblastoma specimens were reviewed by a board-certified pediatric pathologist (F.K.H.), and regions of fetal, embryonal, and mesenchymal histology as well as normal liver were marked. Smart-3SEQ was performed as previously described^33^. Briefly, fresh sections of each FFPE block were cut on a microtome at 7 μm thickness and mounted on glass slides with polyethylene naphthalate membranes (Thermo Fisher Scientific, LCM0522) and stored in a nitrogen chamber. In preparation for laser capture microdissection, slides were immersed sequentially for 20s each in xylenes (three times), 100% ethanol (three times), 95% ethanol (two times), 70% ethanol (two times), water, hematoxylin (Dako S3309), water, bluing reagent (Thermo Fisher Scientific 7301), water, 70% ethanol (two times), 95% ethanol (two times), 100% ethanol (three times), and xylenes (three times). Immediately after staining, cells were dissected on an ArcturusXT LCM System using the ultraviolet (UV) laser to cut the sample and the infrared laser to adhere it to a CapSure HS LCM Cap (Thermo Fisher Scientific, LCM0215). Roughly 500 cells were captured by area, according to density estimates by cell counting on small areas. After LCM, the cap was sealed in a 0.5 mL tube (Thermo Fisher Scientific, N8010611) and stored at −80°C until library preparation. Sequencing libraries were prepared using the Smart-3SEQ protocol (https://genome.cshlp.org/content/suppl/2019/10/24/gr.234807.118.DC1/Supplemental_ <u>File_2.pdf</u>) for FFPE tissue on an Arcturus LCM HS cap, with the pre-SPRI pooling option and a single batch for all samples. Libraries were characterized immediately and stored at −20°C until sequencing. Sequencing was performed on the pooled library using the NextSeq 500/550 High Output V2 kit with 75 cycles (read 1: 76 cycles, index 1: 2bp, index 2: 8bp). Reads were aligned to the human reference genome with STAR and read counts were obtained using featureCounts (Subread). PCA analysis was performed on results from 72 samples (excluding 2 outliers) using FactoMiner on the top 1500 expressed genes. Clustering was performed with Java TreeView using the following parameters: filtering for SD > 2.5 (1222 genes), Pearson correlation (uncentered), complete linkage. Differential expression analyses were performed with DESeq2^34^, excluding 2 outlier samples and collapsing repeated samples.

### Tissue processing

Fresh tumor and normal liver tissue were fixed overnight at room temperature in 10% neutral buffered formalin, dehydrated, cleared with HistoClear (Natural Diagnostics), and embedded in paraffin. Sections were cut at 5 μm thickness, rehydrated, and processed for hematoxylin/eosin staining, immunofluorescence, or in situ hybridization as described below.

### Immunofluorescence

Following deparaffinization and hydration, antigen retrieval of FFPE tissue sections was performed with Tris buffer at pH 8.0 (Vector Labs H-3301) in a pressure cooker for 20 minutes. Slides were permeabilized with phosphate buffered saline (PBS) containing 0.1% Tween-20, then blocked in 5% normal donkey serum in PBS containing 0.1% Triton-X. Sections were incubated sequentially with primary and secondary antibodies and mounted in Prolong Gold with DAPI (Invitrogen). The following antibodies were used: HNF4α (rabbit, 1:50; Santa Cruz sc8987), CK19 (rabbit, 1:100, Abbomax 602-670), Ki-67 (rat, 1:100; eBioscience 14-5698-82), phospho-histone H3 (ser10) (rabbit, 1:1000, Millipore 06-570), activated β-catenin (mouse FITC-conjugated, 1:50, BD β-Catenin clone #14, custom AB #624044; detected with secondary antibody to FITC). Images were captured on a Zeiss Axio Imager Z.2.

### RNAscope in situ hybridization

Single and dual in situ hybridizations were performed using the manual RNAscope 2.5 HD Assay-Red Kit and RNAscope 2.5 HD Duplex Assay Kit (Advanced Cell Diagnostics), respectively, according to the manufacturer’s instructions. Images were taken at 20x magnification on a Zeiss Axio Imager Z.2. Probes used in this study were Axin2 (target region: 502 – 1674), FGF19 (target region: 457 – 2,128), KLB (target region: 729 – 1,680), SOX4 (target region: 5 – 3,238).

### Culture of hepatoblastoma tumoroids

Tumor tissue was collected from patients with hepatoblastoma at the time of biopsy, tumor resection, or liver transplant and processed as previously described^45^. Briefly, tissue was placed immediately in cold Advanced DMEM/F12 with 1x GlutaMAX (Gibco 35050061), 10mM HEPES, and 100U/mL Penicillin/Streptomycin (Gibco 15140122) and kept at 4°C. Tissue was minced with a razor blade, washed with DMEM (high glucose, with GlutaMAX), containing 1% FBS (Omega Scientific Inc. FB-01) and 100U/mL Penicillin/Streptomycin (Gibco 15140122) and centrifuged at 150g x 5 min. Cell clusters were embedded in Matrigel (Growth Factor Reduced, Phenol Red-free, LDEV-free, Corning 356231) and layered with media comprised of 100U/mL Penicillin/Streptomycin (Gibco 15140122), 1x GlutaMAX (Gibco 35050061), 10mM HEPES, 1x B27 supplement (Gibco 17504044), 1x N2 supplement (Gibco 17502048), 1.25mM N-acetyl cysteine (Sigma-Aldrich A9165), 10mM nicotinamide (Sigma-Aldrich N3376), 10mM Y27632 (PeproTech 1293823), 0.1mg/mL Normocin (Invivogen ant-nr-2), in Advanced DMEM/F12 (Gibco 12634010, with glucose, non-essential amino acids, sodium pyruvate, and phenol red). Tumoroids were passaged between 1:4 to 1:10 every 1-2 weeks. Cell clusters were released from Matrigel by incubation in dispase 5U/mL (Stem Cell Technologies 07913) for up to 30 minutes at 37°C, washed twice with liver perfusion media containing 5% FBS, washed with William’s media containing 5% FBS, and re-embedded in Matrigel with tumoroid media as above.

### Single cell RNA sequencing

Tumoroids at P1 or P2 were released from Matrigel using dispase 5U/mL (Stem Cell Technologies 07913), followed by dissociation to single cells with TrypLE Express (Gibco 12605010) and resuspension in William’s media with 10% FBS. Single cell RNA sequencing was performed using the 10x Chromium V2 (HB1) or V3 systems, targeting 5000 cells for each sample. Libraries for HB1, HB4, HB6, and HB7 were sequenced with the Illumina NovaSeq 6000 instrument with NovaSeq S1 v.1.5 Reagent Kits with the following reads: 28 bases Read 1 (cell barcode and unique molecular identifier [UMI]), 8 bases i7 Index 1 (sample index), and 91 bases Read 2 (transcript). Libraries for HB12, HB13, HB14, HB15, HB16, HB17 were sequenced with the Illumina NextSeq 550 High Output instrument with the following reads: 28 bases Read 1 (cell barcode and unique molecular identifier [UMI]), 10 bases i7 Index 1 (sample index), 10 bases i5 Index 2 (sample index), and 90 bases Read 2 (transcript). Sample demultiplexing, barcode processing, single-cell counting, and reference genome mapping were performed using the Cell Ranger Software (v3.0.2, GRCh38.3.0.0 ref genome). Dimensionality reduction by principal component analysis (PCA), graph-based clustering, and UMAP visualization were performed using Seurat (version 4.3.0, R package). In brief, data was filtered to include only genes detected in > 3 cells, cells with > 200 and < 10,000 detected genes, and cells with <15% mitochondrial genes. Data from 10 samples were normalized, combined, and integrated using canonical correlation analysis (CCA) to identify anchors. Integrated data was scaled, regressing out the read depth, percent mitochondrial genes, and percent ribosomal genes. Graph-based clustering and UMAP visualization were performed with the first 30 principal components and a resolution of 0.4.

### DNA isolation and CTNNB1 sequencing

DNA was extracted using standard phenol/chloroform or the Qiagen AllPrep DNA/RNA Kit (Qiagen, Hilden, Germany). CTNNB1 PCR for a 1046bp fragment encompassing exon 2 through exon 4 was performed with the following primers: CTNNB1 ex2-F 5′-AGCGTGGACAATGGCTACTCAA CTNNB1 ex4-R: 5′-ACCTGGTCCTCGTCATTTAGCAGTPCR was performed with an annealing temperature of 70°C, extension time of 1 minute, and 35 cycles. Products were isolated by gel extraction and analyzed by Sanger sequencing with the same primers above.

### RNA isolation and qRT-PCR

Total RNA was extracted using the Qiagen RNAeasy Mini Kit (Qiagen, Hilden, Germany) and reverse transcribed (High Capacity cDNA Reverse Transcription Kit; Life Technologies, Carlsbad, CA) according to the manufacturer’s protocol. Quantitative RT-PCR was performed with TaqMan Gene Expression Assays (Applied Biosystem, Waltham, MA) on an StepOnePlus Real-Time PCR System (Applied Biosystems, Waltham, MA). Relative target gene expression levels were calculated using the delta-delta CT method. Gene Expression Assays used were: GAPDH (Hs99999905_m1, FAM-MGB), AXIN2 (Hs00610344_m1, FAM-MGB), CTNNB1 (Hs00355045_m1, FAM-MGB), DKK1 (Hs00183740_m1, FAM-MGB), TBX3 (Hs00195612_m1, FAM-MGB), FGF19 (Hs00192780_m1, FAM-MGB), FGF8 (Hs00171832_m1, FAM-MGB), FGF9 (Hs00181829_m1, FAM-MGB), KRT19 (Hs01051611_gH, FAM-MGB), HNF4A (Hs00230853_m1, FAM-MGB), SOX4 (Hs00268388_s1, FAM-MGB), all from Thermo Fisher Scientific (Waltham, MA).

### Protein extraction and immunoblotting

Cell pellets were incubated for 15 minutes in ice-cold RIPA buffer (50mM Tris pH 8.0, 150mM NaCl, 1% NP-40, 0.1% SDS, 0.5% Na deoxycholate) with freshly added protease inhibitors (Complete mini, EDTA-free, Roche), phosphatase inhibitors (PhosSTOP, Roche), 1mM PMSF, and 1 mM DTT, at a concentration of 10^7 cells/mL. Lysates were centrifuged at max speed for 15 minutes at 4°C and the protein-containing supernatant was stored at −80°C until use. Protein concentration was measured using the DC Protein Assay (Bio-Rad). Equal amounts (20 μg) were diluted 1:1 in 2x Laemmli buffer, boiled, and loaded for SDS-PAGE (miniPROTEAN TGX pre-cast gel, Bio-Rad). After transfer to a nitrocellulose membrane and blocking in 5% milk/PBST, the following antibodies (in 5% milk/PBST) were used: FGF19 (mouse monoclonal, 1 μg/ml, R&D MAB969), GFP (rabbit polyclonal, 1:5000, OriGene TA150122). Membranes were washed, incubated with the appropriate secondary antibody conjugated with horse radish peroxidase (HRP) and diluted in 5% milk/PBST, and detected by chemiluminescence (Western Lightning Plus ECL, ThermoFisher Scientific).

### Histology and immunostaining of tumoroid cells

Tumoroids were released from Matrigel as described above, fixed in 4% paraformaldehyde at room temperature for 1 hour, washed twice in PBS. Tumoroids were resuspended in Histogel (Thermo Scientific) and distributed into a CHEF plug mold. Plugs containing tumoroids in Histogel were fixed, dehydrated, cleared, and embedded in paraffin identically to primary tissues as above.

Immunofluorescence of hepatoblastoma cells was performed on tumoroid cells that were released from Matrigel using dispase and TrypLE Express and replated in 2D on collagen-coated cover slips. After 24 hours of 2D growth in tumoroid media, cells were fixed with 4% PFA or methanol, washed with PBS. Cells were permeabilized with phosphate buffered saline (PBS) containing 0.1% Tween-20, then blocked in 5% normal donkey serum in PBS containing 0.1% Triton-X, and antibody incubation was performed as for tissue sections described above.

### Colony formation assay

Cells were seeded at a density of 3,000 cells per 20 μL drop of Matrigel and 8 drops per well of a 6-well plate. Media was changed every 3-4 days and colonies were assessed after 2 weeks. After removal of media, colonies in Matrigel drops were incubated for 1 hour with Hoechst 33342 (10 μg/mL) in dispase (5U/mL). Colonies were released from Matrigel by gentle pipetting, washed once with liver perfusion media with 5% FBS, resuspended in the same media and transferred to a 96-well plate (Corning) for imaging. Colonies were imaged with the CellInsight CX7 High-Content Screening (HCS) Platform (Thermo Fisher Scientific) using bright-field, confocal Z-stack, and widefield channels, and quantified using the HCS Studio Cell Analysis Software (Thermo Fisher). Cell debris or dead cells were excluded based on colony size, nuclear intensity, and length-to-width ratio, with the same threshold applied for colony counting in all experiments.

### Cell cycle synchronization and flow cytometry of tumoroid cells

Tumoroids were incubated in media with no growth factors and U0126 5 μM for 24 hours. Media was removed and tumoroids were washed twice with media without growth factors or drug and replaced with either media containing growth factors or media without growth factors. After 35 hours, EdU 10 μM was added for 1 hour, then tumoroids were isolated from Matrigel using dispase and further dissociated to single cells using TrypLE. Single cells were washed with liver perfusion media with 5% FBS and fixed with 70% ethanol. Cells were stained with the Click-iT Plus EdU Alexa Fluor 647 Flow Cytometry Assay Kit (Thermo Fisher C10340) and propidium iodide for DNA content. Flow cytometry was performed on a BD FACS Aria II machine.

### Lentiviral transduction of tumoroids

At 24 hours prior to transfection, 293T cells were plated in 6 well plates at 8 x 10^5 cells / well with DMEM / 10% FBS / NEAA / Pen-Strep (Gibco). Cells growing at ∼70-80% confluency were transfected with 1 μg of the desired expression plasmid (Sigma Aldrich) along with 2^nd^ generation lentiviral packaging and envelope plasmids, 0.75 μg of psPAX2 and 0.25 μg of pMD2.g (gifts from D. Trono, Addgene #12260, #12259). At 36 hrs post transfection, the media containing lentiviral particles was collected and passed through a 0.45 μm filter. Polybrene was added to a final concentration of 4 μg/mL. The filtered media containing lentivirus was added at a 1:1 dilution to target hepatoblastoma tumoroids suspended in media. A total of 4 infections, each spaced 12 hours apart, were performed. Tumoroids were transferred to antibiotic selection 24 hours after the last infection. For puromycin-selectable vectors, tumoroids were cultured in media containing puromycin 2 μg/ml for 3 days prior to isolation of cells for RNA or protein extraction or seeding for colony assays.

Predesigned shRNA in the puromycin-selectable pLKO.1 vector were obtained from Sigma Aldrich, targeting the following sequences – CTNNB1: 5′-GCTTGGAATGAGACTGCTGAT (TRCN0000003845) FGF19: 5′-GCTTTCTTCCACTCTCTCATT (TRCN0000040260) SOX4: 5′-AGCGACAAGATCCCTTTCATT (TRCN0000018214) Luciferase: 5′-CGCTGAGTACTTCGAAATGTC (SHC007).

Other plasmids used in the study were: pLKO.1-puro-CMV-TurboGFP (Sigma-Aldrich, SHC003), pLenti-C-mGFP-P2A-puro-FGF19 (Origene, RC203750L4), EdTP (dominant negative Tcf4, Addgene #24311).

