## Supplemental Information including Tables and Figures for "Cholangiocytic differentiation drives cell proliferation in hepatoblastoma through Wnt-dependent FGF19 signaling"

### Table S1 (related to Figure 1).
### Characteristics of hepatoblastoma specimens used for Smart-3SEQ

| Pt | Age (months) | Sex | Histology | Prior Chemo | FGF19 | |
|---|---|---|---|---|---|---|
| | | | | | Smart-3SEQ | In situ |
| **1** | 29 | F | Fetal & Embryonal | N | - | |
| **2** | 9 | M | Predominant Embryonal | N | + | n.d. |
| **3** | 9 | F | Mixed | Y | + | - |
| **4** | 27 | M | Mixed | Y | - | |
| **5** | 8 | F | Mixed | Y | + | - |
| **6** | 34 | M | Fetal & Embryonal | N | + | + |
| **7** | 10 | M | Mixed | N | + | + |
| **8** | 24 | M | Fetal & Embryonal | N | + | + |
| **9** | 37 | F | Mixed | Y | - | |
| **10** | 81 | F | Fetal & Embryonal | N | + | + |
| **11** | 5 | F | Fetal & Embryonal | N | + | + |
| **12** | 12 | F | Fetal & Embryonal | N | - | |
| **13** | 8 | M | Fetal & Embryonal | N | - | |
| **14** | 34 | M | Fetal & Embryonal | N | - | |
| **15** | 27 | F | Mixed | Y | - | |
| **16** | 83 | F | Mixed | Y | - | |
| **17** | 21 | M | Mixed | Y | - | |

Characteristics of the patients and tumors reported are from the date of the specimen. n.d. = not determined.

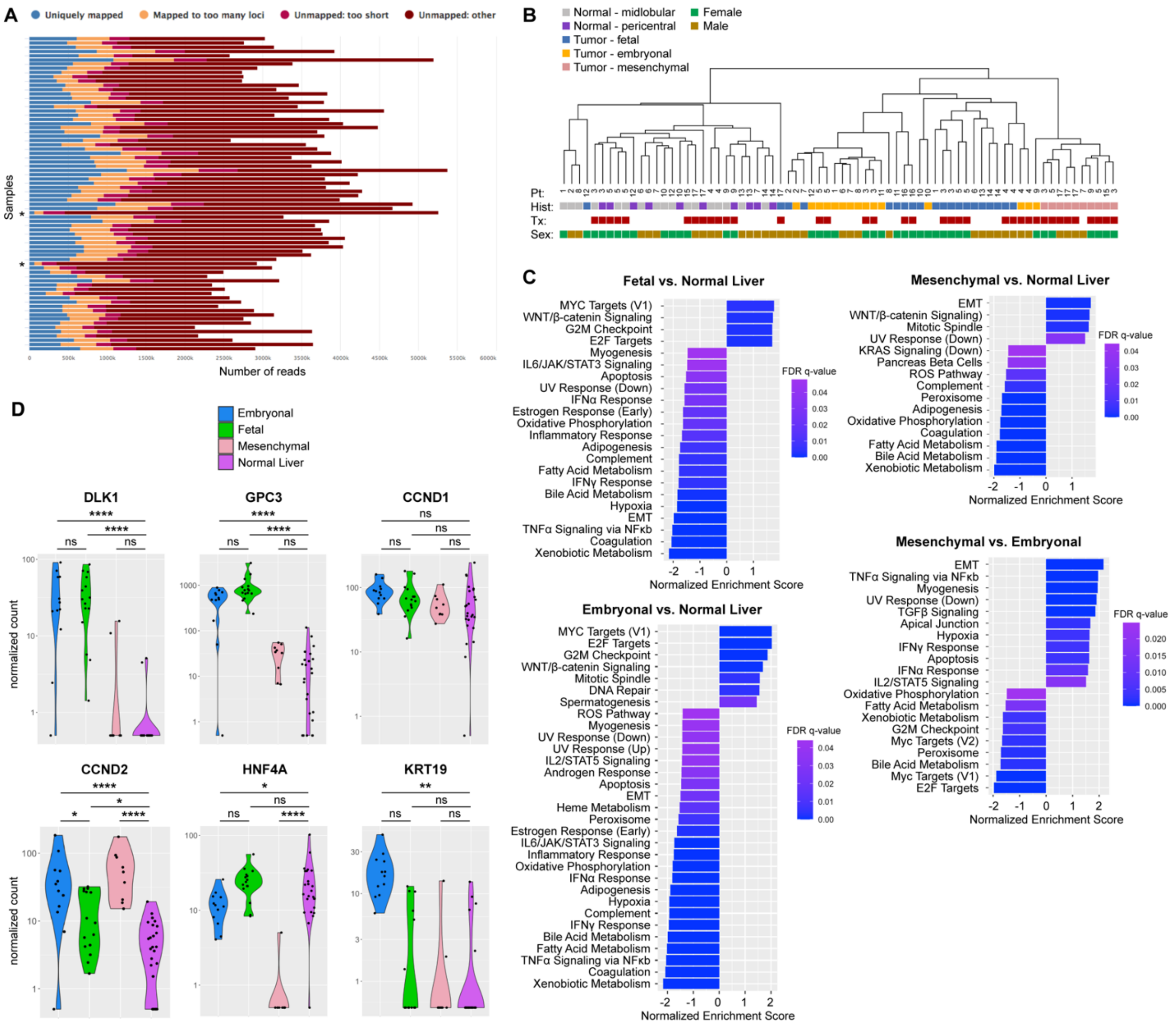


**Figure S1 (related to Figure 1).** A. Read alignment of samples assayed by Smart-3SEQ, using STAR. * indicates samples with poor quality that were excluded from further analysis. B. Unsupervised hierarchical clustering of results from Smart-3SEQ. C. GSEA of differentially expressed genes obtained by DESeq2 comparing Fetal vs. Normal Liver, Embryonal vs. Normal liver, Mesenchymal vs. Normal Liver and Mesenchymal vs. Embryonal regions of hepatoblastoma. D. Normalized expression of DLK1, GPC3, cyclins CCND1 and CCND2, hepatic marker HNF4A, and cholangiocytic marker KRT19 in different regions of hepatoblastoma. * denotes padj <0.05, ** denotes padj <0.01, *** denotes padj <0.001, **** denotes padj <0.0001. ns = not significant.

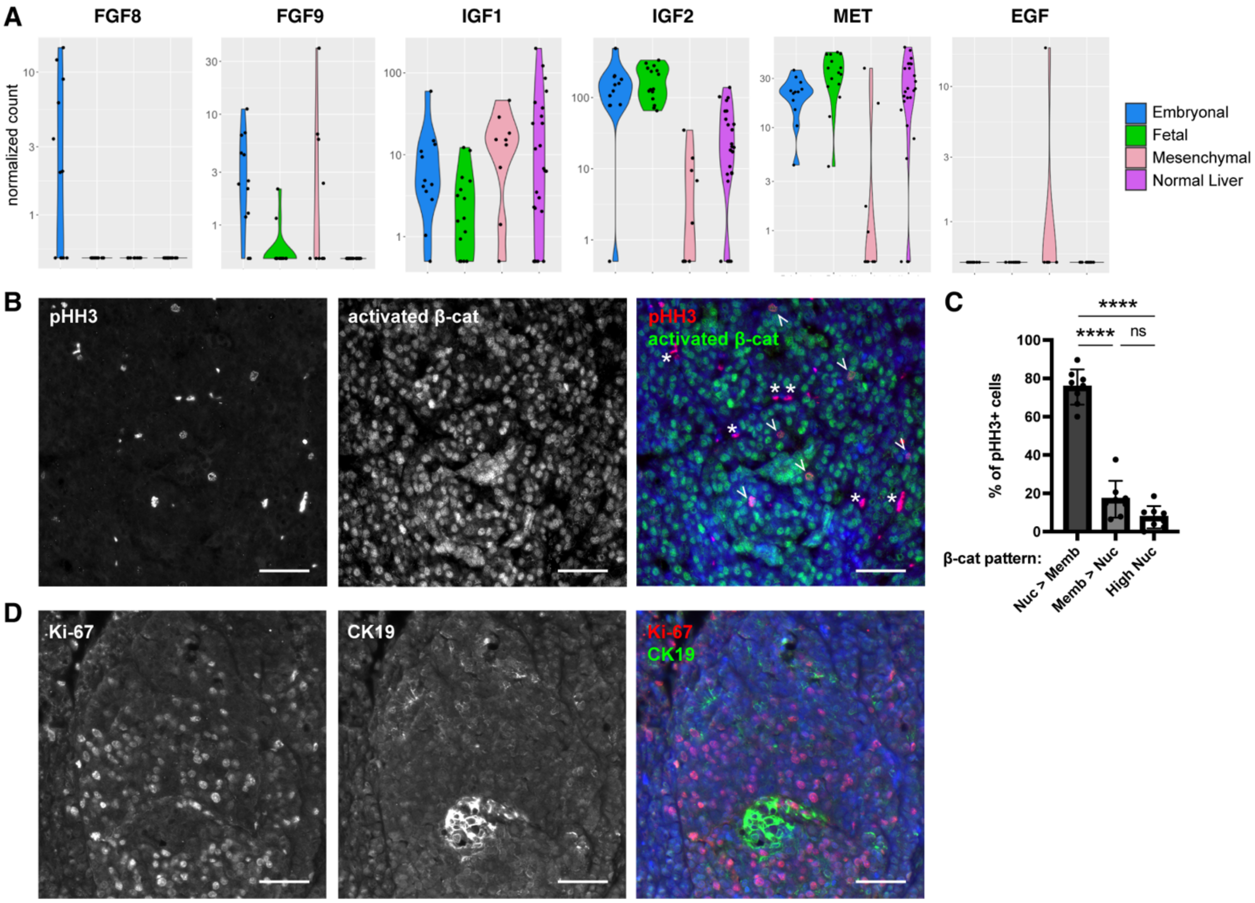


**Figure S2 (related to Figure 2).** A. Normalized expression of FGF8, FGF9, IGF1, IGF2, MET, and EGF in different regions of hepatoblastoma determined by Smart-3SEQ and DESeq2. B. Co-immunofluorescence of phospho-histone H3 (pHH3) (left) and activated β-catenin (middle), merged with DAPI (right). White arrowheads in right panel indicate pHH3+ cells. White asterisks in right panel indicate non-specific staining of blood cells. Scale bar: 50 μm. C. Quantification of percentage of pHH3+ cells with different patterns of β-catenin staining as indicated. **** denotes $p < 0.0001$ by ordinary one-way ANOVA ($n = 8$ independent specimens, >50 pHH3+ cells scored for each experiment). D. Co-immunofluorescence of Ki-67 (left) and CK19 (middle), merged with DAPI (right). Scale bar: 50 μm.

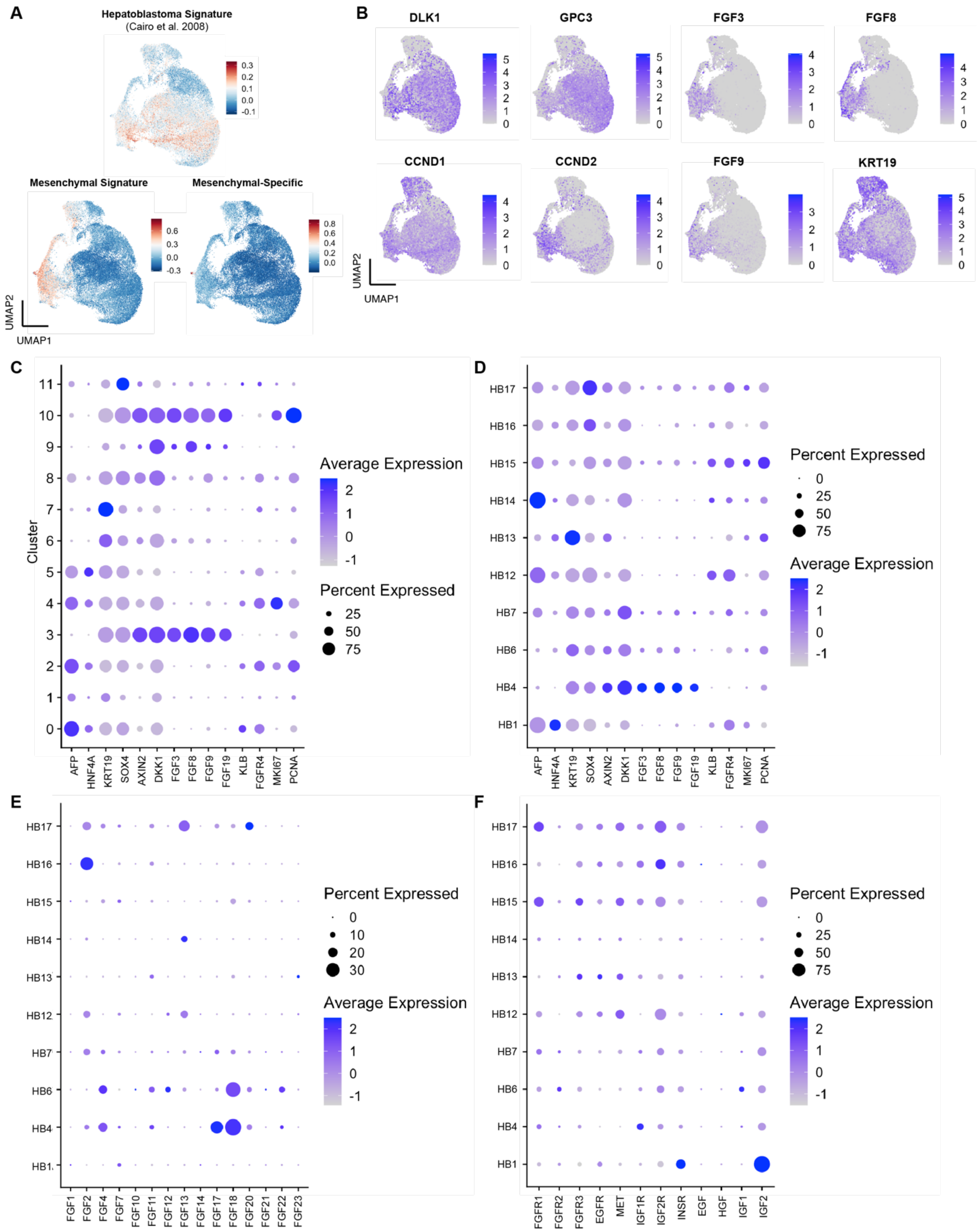


**Figure S3 (related to Figure 3).** A. UMAP plots representing relative expression of the following gene signatures defined by lists in Table S6: hepatoblastoma signature[1], mesenchymal gene signature defined as the top 50 differentially upregulated genes between the mesenchymal histology and normal liver, and mesenchymal-specific gene signature defined as the top 50 differentially upregulated genes between the mesenchymal and embryonal histologies, identified by Smart-3SEQ and DESeq2 as in Figure 1. B. UMAP plots showing relative gene expression for hepatoblastoma markers, cyclins, and FGFs. C. Dot plot showing relative gene expression of markers in different clusters. D. Dot plot showing relative gene expression of markers in different HB tumoroids. E. Dot plot showing relative expression of all FGFs in different HB tumoroids. F. Dot plot showing relative expression of all FGFRs and other growth factor/receptor combinations in different HB tumoroids.

#### Table S4 (related to Figure 3). Characteristics of hepatoblastoma tumoroids

| | Age (mo) | Sex | Histology | Stage/Risk | Prior Chemotherapy | Specimen source | CTNNB1 mutation | Other mutations (VAF) | Status |
|---|---|---|---|---|---|---|---|---|---|
| **HB1** | 129 | M | Fetal & Embryonal | Relapsed/metastatic | Y | Lung metastasis | A21_G38del | n.d. | Deceased |
| **HB4** | 29 | M | Mixed (with teratoid) | Intermediate Risk (later relapsed) | Y | Resection | A5_S37del | FGF4 P175S (48%)<br>SMAD4 I525V (50%) | CR2 |
| **HB6** | 42 | M | Mixed (with teratoid) | Relapsed | Y | Resection | D32N | AR A646 (99%)<br>FGFR2 c1287+2A>G (38%) | Deceased |
| **HB7*** | 21 | unknown | Fetal & Embryonal | Unknown | Y | Resection | S23_S33delinsP | n.d. | unknown |
| **HB12** | 15 | F | Fetal & Embryonal | Low Risk | N | Biopsy | G34V | ARID1A W1844ter (germline)<br>otherwise n.d. | CR1 |
| **HB13** | 9 | M | Mixed (with teratoid) | Intermediate Risk | Y | Resection | G34R | APC I2341X (germline)<br>FGF4 A29V (48%) | CR1 |
| **HB14** | 4 | F | Mixed (with teratoid) | Intermediate Risk | N | Biopsy | A5_D63del | GATA3 A396T (47%) | CR1 |
| **HB15** | 34 | M | Mixed (with teratoid) | Low Risk (later relapsed) | Y | Resection | A5_D63del | TSC2 V1034I (52%)<br>APC N2109H (49%)<br>ATM H231R (47%) | CR2 |
| **HB16** | 24 | F | Fetal & Embryonal | High Risk (metastatic) | N | Biopsy | A5_D63del | LRP1B E71* (58%)<br>BCR V1107I (18%)<br>CDKN1B L120M (43%)<br>SMARCA4 P913L<br>BCR T1018A (14%)<br>NFE2L2 E79_E82del (25%) | Ongoing treatment |
| **HB17** | 7 | F | Mixed | Intermediate Risk | N | Resection | A5-D63del | ARID1A N106fs (13%)<br>BTK R492H (50%)<br>ERBB2 P625L (46%)<br>KDR A772V (45%)<br>FOXR2 S92N (41%)<br>FGF4 A24T (35%) | CR1 |

Characteristics of the patients and tumors reported are from the date of the specimen. n.d. = not determined.
*Gift from Bruce Wang[2]

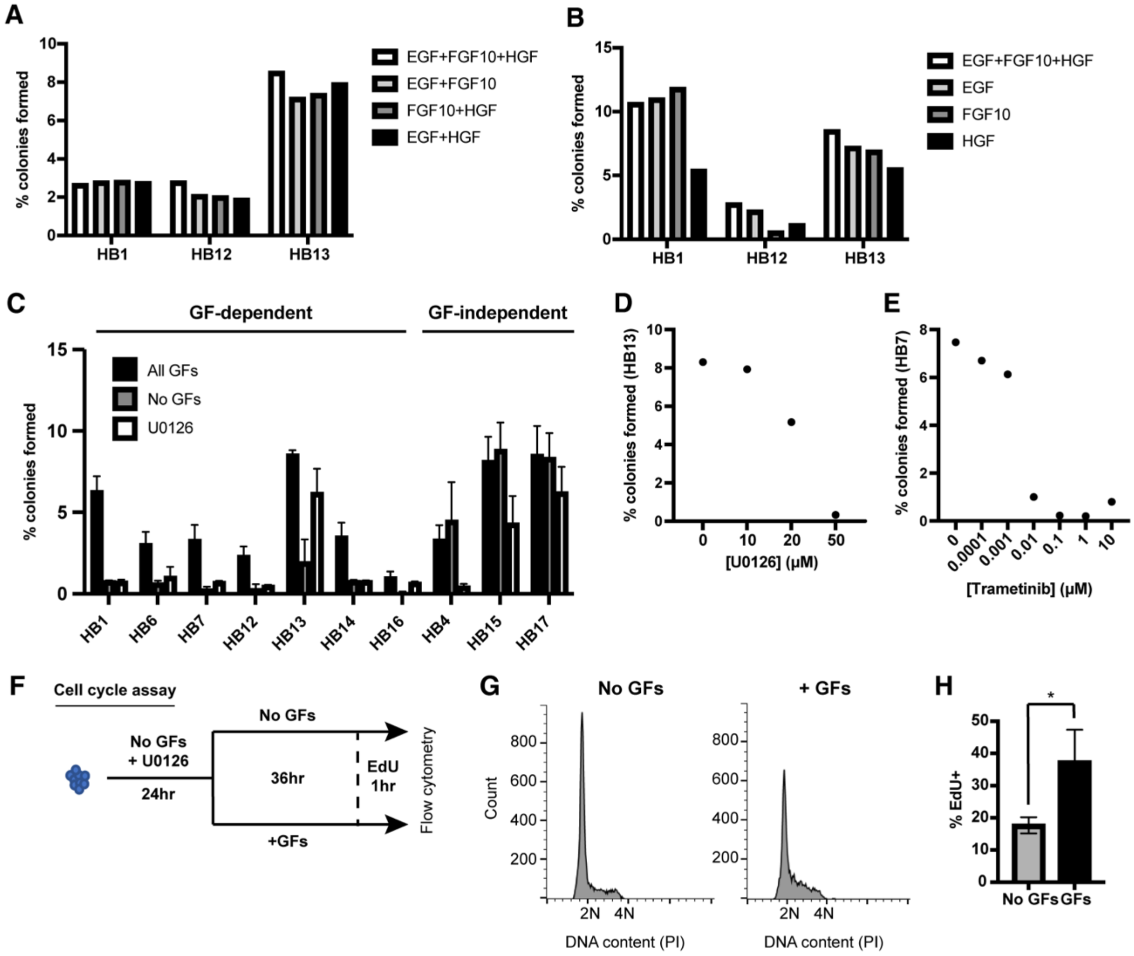


**Figure S4 (related to Figure 4).** A. Colony formation assay of single cells from the indicated tumoroids growing in media containing all GFs (EGF 50 ng/ml + FGF10 100 ng/ml + HGF 25 ng/ml), EGF 50 ng/ml + FGF10 100 ng/ml, FGF10 100 ng/ml + HGF 25 ng/ml, or EGF 50 ng/ml + HGF 25 ng/ml. B. Colony formation assay of single cells from the indicated tumoroids growing in media containing all GFs (EGF 50 ng/ml + FGF10 100 ng/ml + HGF 25 ng/ml), EGF (50 ng/ml), FGF10 (100 ng/ml), or HGF (25 ng/ml). C. Quantification of colony assay in media with all GFs, no GFs, or all GFs + U0126 (5 μM) for each of 10 tumoroids (data representing All GFs and No GFs are the same as in Fig. 4B, repeated here for comparison to the drug treated condition). D. Colony formation of HB13 in media contain all GFs and U0126 at the different concentrations indicated. E. Colony formation of HB7 in media contain all GFs and trametinib at the different concentrations indicated. F. Schema for cell cycle assay in G and H. G. Representative flow cytometry analysis of HB1 cells growing in media with no GFs and with U0126 (5 μM) for 24hrs, then released into media with no GFs or with all GFs for 36 hrs with 1 hr incubation in EdU prior to harvest, stained with PI for DNA content. H. Quantification of %EdU+ cells from flow cytometry experiment in D (n = 3 independent experiments, error bars indicate standard deviation. * denotes $p<0.05$ by paired student t-test).

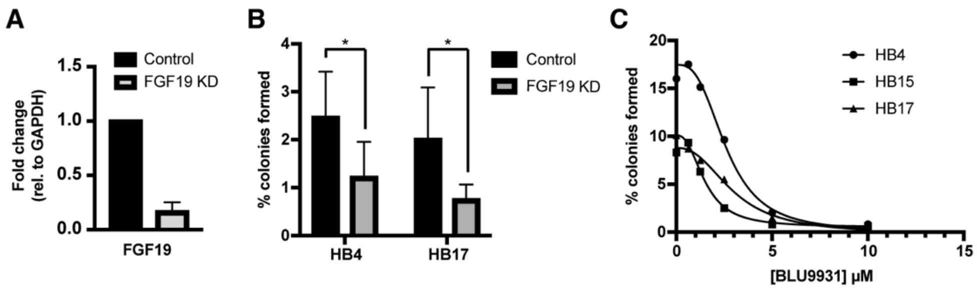


**Figure S5 (related to Figure 5).** A. qRT-PCR for FGF19 in HB15 cells transduced with lentiviral shRNA to luciferase (control) or FGF19, assayed after 3 days of antibiotic selection (n = 3 independent experiments, error bars indicate standard deviation). B. Quantification of colony assay for HB4 and HB17 in the setting of control shRNA (to luciferase) vs. FGF19 shRNA (n = 3 independent experiments, error bars indicate standard deviation. * denotes $p<0.05$ by paired student t-test). C. Colony formation assay of single cells from the indicated tumoroids in media containing all GFs and FGFR4 inhibitor, BLU9931, at the concentrations indicated.

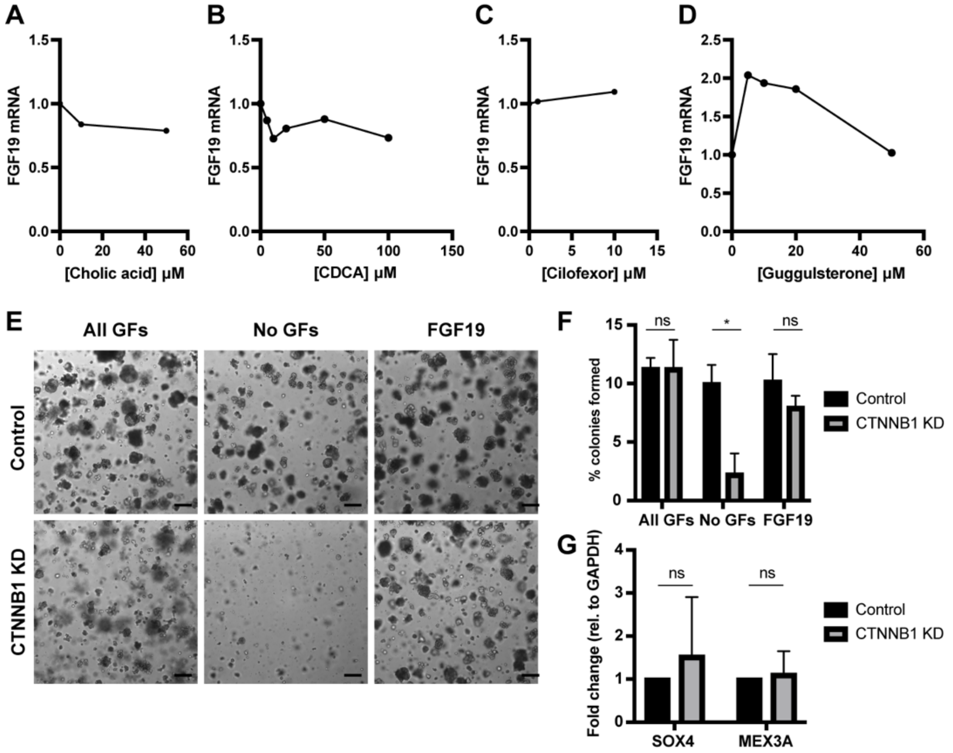


**Figure S6 (related to Figure 6).** A. qRT-PCR for FGF19 (relative to GAPDH) in HB15 cells grown in media with all GFs after 24 hour incubation with cholic acid at the concentrations indicated. B. qRT-PCR for FGF19 (relative to GAPDH) in HB15 cells grown in media with all GFs after 24 hour incubation with chenodeoxycholic acid at the concentrations indicated. C. qRT-PCR for FGF19 in HB15 cells grown in media with all GFs after 48 hour incubation with FXR agonist, cilofexor, at the concentrations indicated. D. qRT-PCR for FGF19 in HB15 cells grown in media with all GFs after 48 hour incubation with FXR antagonist, guggulsterone, at the concentrations indicated. E. Brightfield images of HB15 colonies transduced with lentiviral shRNA to luciferase (control) or CTNNB1, seeded as single cells after 3 days of antibiotic selection, and grown for 2 weeks in media containing EGF + FGF10 + HGF (All GFs), no GFs, or FGF19. Scale bar: 200 $\mu$m. F. Quantification of colony assay as in B (n = 3 independent experiments, error bars indicate standard deviation. * denotes $p<0.05$ by paired student t-test, ns = not significant). G. qRT-PCR for SOX4 and MEX3A in HB15 cells transduced with lentiviral shRNA to luciferase (control) or CTNNB1, assayed after 3 days of antibiotic selection (n = 3 independent experiments, error bars indicate standard deviation. ns = not significant by paired student t-test).
